# Chemical proteomics decrypts the kinases that shape the dynamic human phosphoproteome

**DOI:** 10.1101/2025.11.18.689017

**Authors:** Florian P. Bayer, Julian Müller, Nicole Kabella, Miriam Abele, Yun-Chien Chang, Leonardo Daniel Estrada Duenas, Cecilia Jensen, Firas Hamood, Chien-Yun Lee, Annika Schneider, Matthew The, Bernhard Kuster

## Abstract

Mass-spectrometry-based phosphoproteomics enables the analysis of thousands of protein phosphorylation events across the human proteome. However, there is a lack of scalable, hypothesis-free, and statistically sound approaches for discovering, evaluating, and falsifying kinase::substrate relationships (KSRs). Here, we developed a new concept termed potency-coherence analysis. By measuring and integrating 17 million peptidoform-specific dose-response curves for 133 kinase inhibitors with known targets and affinities, we could critically re-evaluate published KSRs and discover thousands of potency-coherent and motif-plausible new KSRs for 96 human kinases. Application of these high-confidence KSRs enabled the estimation of kinase and signaling pathway activities in cancer patient biopsies. This unified and extendable framework has been implemented in ProteomicsDB to aid researchers in understanding the human phosphoproteome in health and disease.

## Introduction

Eukaryotic cells have evolved a remarkably versatile system of reversible protein post-translational modifications (PTMs), which enables rapid modulation of protein structure and function in response to changing environments (*1, 2*). Protein phosphorylation stands out in that the human genome encodes 518 kinases that phosphorylate proteins at serine (pS), threonine (pT), and tyrosine (pY) residues, and 189 phosphatases that remove these PTMs (*3, 4*). Recent estimates suggest that most human proteins can be phosphorylated at multiple positions (*5*) and that 1-6 million molecularly distinct such proteoforms may exist (*6*). While phosphorylation is central to processes such as signal transduction, the functions and kinase::substrate relationships (KSRs) of most phosphorylation sites are currently unknown (*7*). This knowledge gap hampers our understanding of proteoform-based diseases like cancer, which are often caused or accompanied by aberrant kinase/phosphatase activity (*8–10*).

A systematic understanding of the KSRs that shape the human phosphoproteome faces multiple practical challenges. First, the current reliance on a limited set of antibodies targeting a few well-characterized sites narrows the perspective to a small area of biology, and ongoing concerns regarding antibody specificity question many reported findings (*11, 12*). Second, kinase inhibitors, when used as research tools to study PTMs, are often applied at too high concentrations or for too long. This complicates data interpretation because i) inhibitors often lack selectivity and bind on- and off-targets with different potencies (*13*), ii) the effect of kinase inhibition propagates through cell type-specific signaling pathways, indirectly altering additional kinase activities, and iii) protein expression or cell population changes resulting from extended treatment times can further obscure causes and consequences (*14*). Together, these issues impede attributing a change in phosphorylation to the activity change of a particular kinase. Third, public databases cannot curate data at the scale produced today. The concatenation of data from high-throughput studies of varying scientific rigor has led to an alarming growth in false-positive annotations that outpaces the addition of true-positive cases (*15*). Yet, computational and manual analyses depend heavily on these resources (*16, 17*). As a result, little to nothing is known about most KSRs despite the wealth of existing data, and distinguishing true from false KSRs is becoming increasingly difficult.

Therefore, there is a pressing need to rigorously identify the kinases that shape the human phosphoproteome. Recent progress includes the Kinase Library (*18, 19*), a computational tool developed on the basis of large-scale in vitro kinase assays that refined substrate phosphorylation motifs for almost all human kinases. Although this does not yield direct KSRs, it narrows the list of candidate kinases for any phosphorylation site of interest. We recently developed dose-dependent PTM profiling (decryptM) to elucidate the cellular mechanisms of action(s) (MoA) of small molecule inhibitors (*20–22*). Here, we turned the concept around to elucidate KSRs by large-scale phosphoproteome perturbation experiments using 133 kinase inhibitors with known targets and binding affinities in five cancer cell lines. To integrate the millions of measured quantitative, peptidoform-specific dose-response profiles, we developed a unified statistical framework, termed “potency coherence analysis”, and implemented it in ProteomicsDB (*23*). This new concept enabled a critical re-evaluation of published KSRs and the discovery of thousands of confident new KSRs for 96 human kinases. We highlight the translational potential of the approach by estimating kinase activities in thousands of cancer patient phosphoproteomes with potential therapeutic utility. When combined with ongoing chemical biology efforts to create chemical probes for every kinase (*24, 25*), our strategy can be expanded to cover all kinases and their substrates, advancing fundamental biology and precision medicine alike.

## Results

### DecryptM profiling at scale describes the landscape of drug-perturbed phosphoproteomes

To capture many kinase activities in different cellular contexts, we optimized a high-throughput phosphoproteomics workflow and systematically measured dose-response characteristics of 133 clinical kinase inhibitors across five cancer cell lines (see materials and methods; fig. 1A). Because inhibitors bind to on- and off-target kinases with specific affinities, observed phosphorylation changes were a function of the applied dose. Regression of dose-response models yielded effective concentration (pEC_50_=-log_10_ EC_50_) and effect size (log_2_ fold change) estimates for each phosphorylated peptide (peptidoform). Curves were classified as up-, down-, not-, or unclearly regulated using CurveCurator’s relevance score (*26*). It integrates statistical significance and biological effect size, and thereby enables an objective assessment of the presence or absence of drug regulation at low error rates. All MS raw files and dose-response curves are publicly available once approved (*27, 28*), but they can be interactively explored in ProteomicsDB (proteomicsdb.org/decryptm) already.

**Fig. 1.**
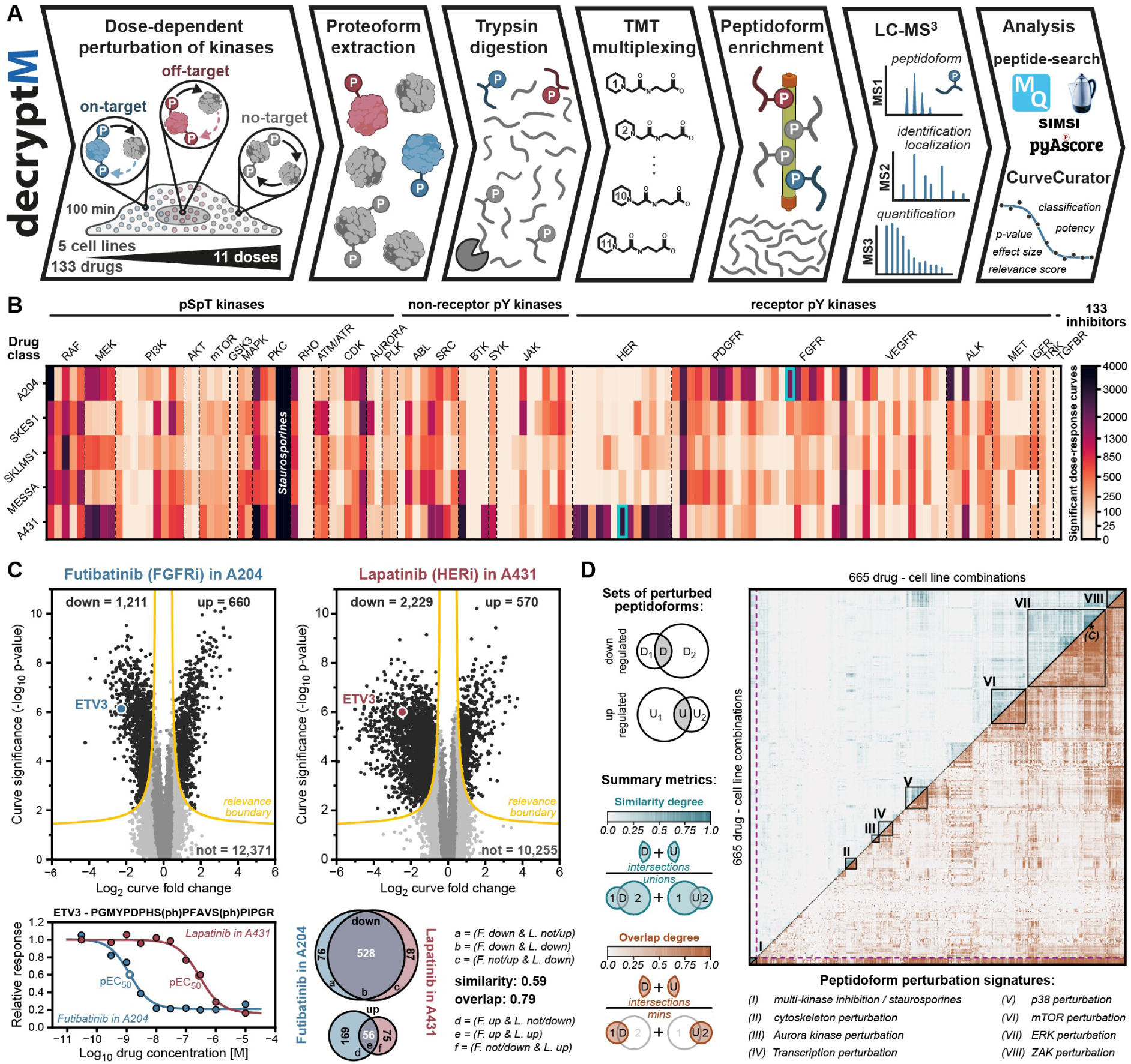
DecryptM analysis of context-specific drug-perturbed phospho- proteomes at scale. ***(A)*** *Schematic representation of the decryptM workflow used to measure 17 million dose- dependent phosphorylated peptidoform profiles in response to 133 kinase inhibitors in 5 cancer cell lines. **(B)** Heatmap summarizing the number of significant drug-perturbed peptidoforms across 665 decryptM experiments. Drugs were arranged by drug target class. Futibatinib (FGFRi) in A204 and Lapatinib (HERi) in A431 are highlighted with a turquoise box.* ***(C)*** *top panel: volcano plot summaries of the decryptM experiments marked in B. Each dot is a peptidoform dose-response curve. Colors denote different CurveCurator classifications (black: “up-” or “down-regulated”; dark gray: “not-regulated”, light gray: “unclear”). The yellow line represents CurveCurator’s relevance boundary (alpha=0.05, fc_lim_=0.45). Bottom right panel: Dose-response curves of one peptidoform of the transcription factor ETV3 highlighted in the volcano plots. Bottom right panel: Venn diagrams depicting the overlap of up- or down- regulated peptidoforms between both experiments. **(D)** Heatmap showing the degree of similarity (blue, top left triangle) and overlap (brown, bottom right triangle) between any pair of decryptM combinations. Hierarchical clustering was based on the degree of overlap. Prominent clusters are labeled by roman numerals and represent shared peptidoform perturbation signatures of common signaling axes. The decryptM comparison from panel C is indicated with an asterisk. The mathematical calculation of similarity and overlap is iconized on the left*.

The complete 665-experiment matrix (5 cell lines × 133 inhibitors; fig. 1B) comprised 7,315 perturbations (665 decryptM experiments with 11 doses). On average, ∼24,000 peptidoforms were profiled per drug/cell-line experiment (total 90,776 peptidoforms), resulting in ∼17.1 million peptidoform dose-response curves (fig. S1). The 100-minute treatment minimized confounding contributions from protein expression changes or cell population shifts (*14, 29*), but also restricted the analysis to “dynamic” phosphorylation events with active kinase-phosphatase pairs. In the absence of counteracting phosphatase activity, phosphorylation sites remained “static” and could not respond within this short timeframe. Globally, most drugs altered the abundance of 1-5% of peptidoforms. Notable exceptions were broad-spectrum inhibitors of the Staurosporine family (Lestaurtinib, K252a), perturbing up to 30%. By contrast, highly selective BTK or JAK inhibitors regulated <20 peptidoforms across cell lines, consistent with absent or inactive kinase targets.

Next, we performed pairwise comparisons of decryptM experiments using sets of up-, down-, and not-regulated peptidoforms to assess the similarity and overlap between decryptM experiments. For example (fig. 1C), the decryptM profile of A204 sarcoma cells treated with the pan-FGFR inhibitor Futibatinib strongly resembled that of EGFR-amplified A431 epithelial cells treated with the dual EGFR/ERBB2 inhibitor Lapatinib, despite non-overlapping targets (*30*) and distinct cancer biologies (*31*). This is because both decryptM profiles were dominated by shared peptidoforms from the ERK signaling axis that is downstream of FGFR in A204 and EGFR in A431 and included e.g. an ERK phosphorylation site on the transcription factor ETV3 (*32*), which is modulated by both drugs with different potencies. Extending the analysis to all 665 experiments, we observed clustering by major downstream pathways but not by cell line or designated drug target class (fig. 1D; fig. S2). The overall degree of similarity was very low (mean <0.02). At the same time, >50% of decryptM combinations shared regulated peptidoforms. For instance, the decryptM profile of the multi-kinase inhibitor Lestaurtinib almost completely contained the profiles of Trametinib (selective MEKi), Losmapimod (p38i), Temsirolimus (MTORi), and Dinaciclib (panCDKi) (fig. S2D). These findings indicate that i) the 133 inhibitors elicit diverse kinase activity changes, and ii) drug-perturbed sets of peptidoforms are highly context-dependent both within and across cell lines. Such context dependencies result from the presence and activity of specific kinases and phosphatases, their integration into shared or distinct signaling pathways, and whether inhibitors share on- or off-targets. Therefore, the simple idea of drug or kinase “signatures” (*33–37*), which are derived from individual or multiple one-dose-perturbed sets of peptidoforms, cannot by themselves resolve causal links between dynamic peptidoform changes and the underlying activity of specific kinases.

### The potency dimension links biologically connected peptidoforms

As long as phosphatase activity is not limiting, peptidoforms change abundance at a pEC_50_ that corresponds to the drug’s binding affinity to its target kinase(s), defined by the dissociation constant K_D_ (fig. 2A). Consequently, all dynamic substrates of an inhibited kinase or any downstream kinase should respond with similar pEC_50_. Because inhibitors usually target multiple kinases with different affinities, the perturbed phosphoproteome is a mixture of peptidoforms of different but distinct pEC_50_. Below and in the following paragraphs, we show that the potency with which peptidoforms respond to kinase activity changes - regardless of their position in a signaling cascade - is the key dimension for deconvoluting drug-perturbed phosphoproteomes into the responsible kinases. For the sake of simplicity, these concepts are illustrated using the same example of the EGFR inhibitor Afatinib in the EGFR-dependent cell line A431.

**Fig. 2.**
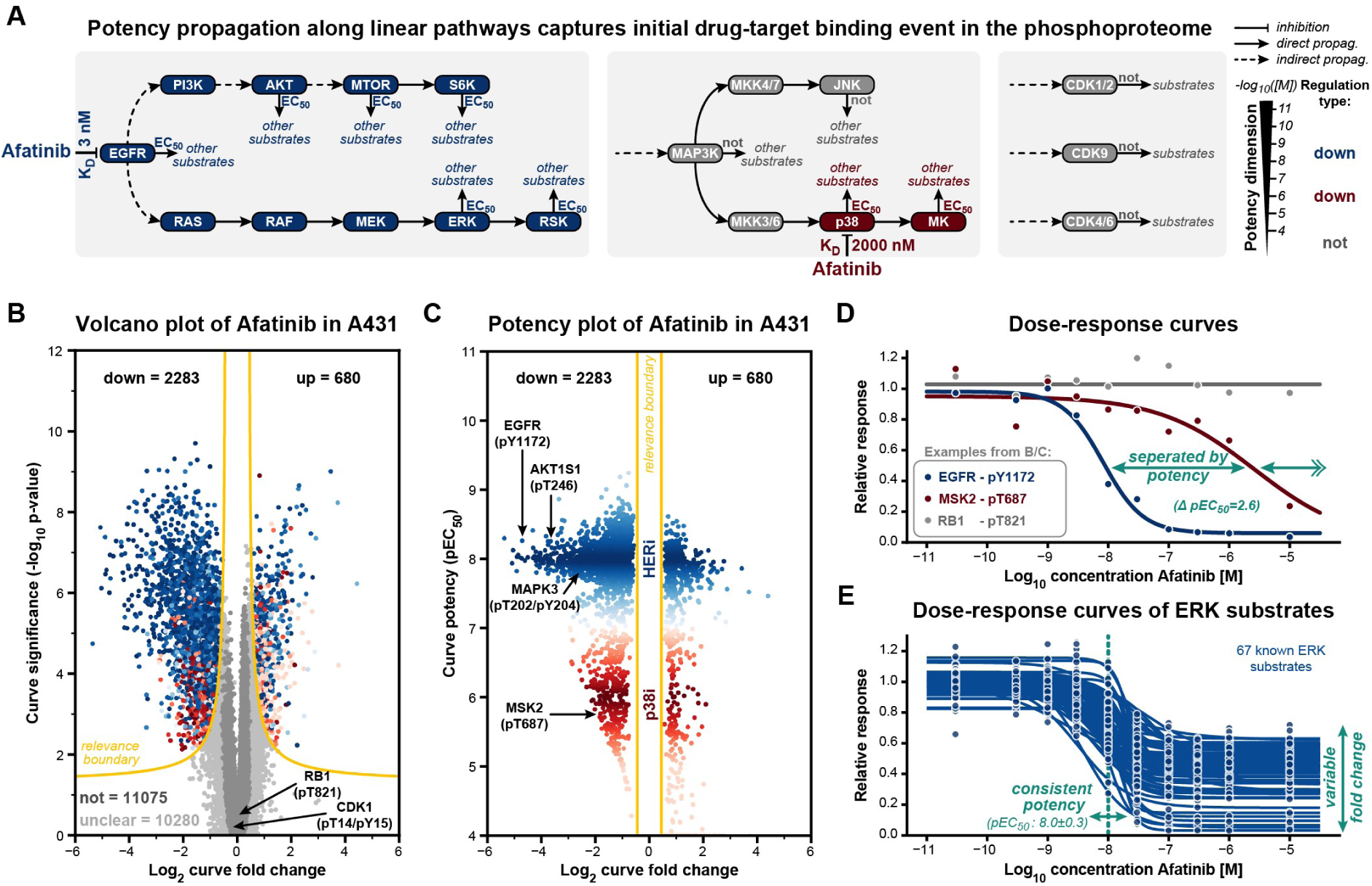
The potency dimension distinguishes signaling pathways engaged by drug targets of different binding affinities. ***(A)*** *Signaling pathway schematics of A431 cells. Left panel: dose-dependent inhibition of EGFR by Afatinib leads to a dose-dependent activity reduction of all downstream kinases and dose-dependent abundance changes of substrates with an effective drug concentration (EC_50_) that is similar to the dissociation constant (K_D_) of EGFR:Afatinib binding (blue pathway nodes). Middle panel: same as right panel, but for off-target inhibition of p38. Middle and left panel: Kinases that are neither directly inhibited nor indirectly perturbed do not change activity (grey pathway nodes), and their substrates do not change abundance. **(B)** CurveCurator volcano- plot for Afatinib in A431, akin to* fig. 1C*, but coloring of regulated peptidoforms by potency (blue: very potent, red: weakly potent). **(C)** CurveCurator potency plot for Afatinib in A431 shows two populations of regulated peptidoforms separated by potency. Blue cases (very potent) result from EGFR on-target inhibition and red cases (weakly potent) from p38 off-target inhibition. **(D)** Dose-response curves illustrating highly potent, weakly potent, and “not”- regulated peptidoforms from panels B, C. **(E)** Overlay of 67 dose-response curves of known and Afatinib-responsive ERK substrates. While effect sizes vary a lot, potencies are very consistent around 10 nM*.

In this example, 2,283 peptidoforms showed decreased and 680 increased phosphorylation in a dose-dependent manner (fig. 2B). While EGFR is the receptor tyrosine kinase (RTK) driving A431 proliferation and sits at the top of the signaling cascade, only a few regulated peptidoforms are direct EGFR substrates. Instead, the majority arise from downstream kinases. In addition, Afatinib also inhibits p38 as an off-target, contributing to the observed effects. Similar to standard replicated one-dose perturbation experiments, neither the significance nor the effect size of dose-response curves can distinguish which initial kinase inhibition triggered which peptidoform changes, as discussed above (fig. 2B). By contrast, plotting pEC_50_ against effect size revealed a bimodal distribution in the potency dimension (fig. 2C). This is because Afatinib binds and inhibits EGFR with ∼3 nM K_D_ and p38 with ∼2,000 nM K_D_ (*30*). Accordingly, downstream phosphorylation events on MAPK3-pT202/pY204 and AKT1S1-pT246, as well as hundreds of other peptidoforms, match the potency of the auto-phosphorylation site EGFR-pY1172, while MSK2-pT687 phosphorylation tracks p38 inhibition (fig. 2D). Importantly, the many not-regulated substrates of kinases that are present in the data, such as the CDK2/4/6-substrate RB1-pT821 (*38*), provide evidence for the absence of kinase perturbation. Such quantitative distinctions are only possible with dose-response measurements, and we note that high-quality negative data is as important as a responsive pEC_50_ for the following analyses.

Across all 665 drug-cell line combinations, about half displayed multimodal or broadened pEC_50_ distributions reflecting widespread polypharmacology (fig. S3A-C) (*30*). While not always as clear as with Afatinib, the potency dimension consistently linked observed peptidoform pEC_50_ to kinase inhibition even for very unselective inhibitors like Dasatinib (fig. S3D). By contrast, the effect size dimension was much less consistent, as peptidoform fold changes depend on multiple factors, including phosphorylation/dephosphorylation kinetics, structural site accessibility, and protein localization. This is exemplified by 67 annotated ERK substrates in the Afatinib-A431 dataset (fig. 2E). While effect sizes varied widely (1.6 - 30 fold reduction), their potencies were very similar (mean ± std pEC_50_: 8.0 ± 0.3).

### Estimating the potency of kinase activity changes requires high-quality kinase::substrate annotations

Following the logic above, peptidoform dose-response curves are indirect measures of kinase activity. If bona fide, kinase-unique substrate peptidoforms exist and were known, then their averaged pEC_50_ could be used to estimate the drug concentration at which the responsible kinase directly or indirectly changed activity. We term this the “inferred kinase perturbing concentration” (inf_pEC_50_). For functionally closely related kinases with strongly overlapping substrate spaces, e.g. MAP2K1/2 or MAPK1/3, “kinase groups” such as MEK or ERK can be defined to simplify analysis (fig. 3A). The central challenge for systematically inferring the potency of kinase activity changes in this way is to find suitable KSRs. PhosphoSitePlus (PSP) is currently the most comprehensive KSR database (*5*). Of the 90,776 unique peptidoforms in our dataset, ∼47% were regulated in at least one decryptM experiment, yet only ∼8% could be annotated with kinases from PSP (fig. 3B). The annotations were dominated by CDK1/2, MAPK1/3, GSK3B, and AKT1, reflecting a strong general bias towards widely studied kinases in PSP. We note that some kinases, e.g., EEF2K and LIMK1/2, naturally have narrow physiological substrate spaces due to specialized biological functions (*39, 40*), explaining the limited peptidoform matches.

**Fig. 3.**
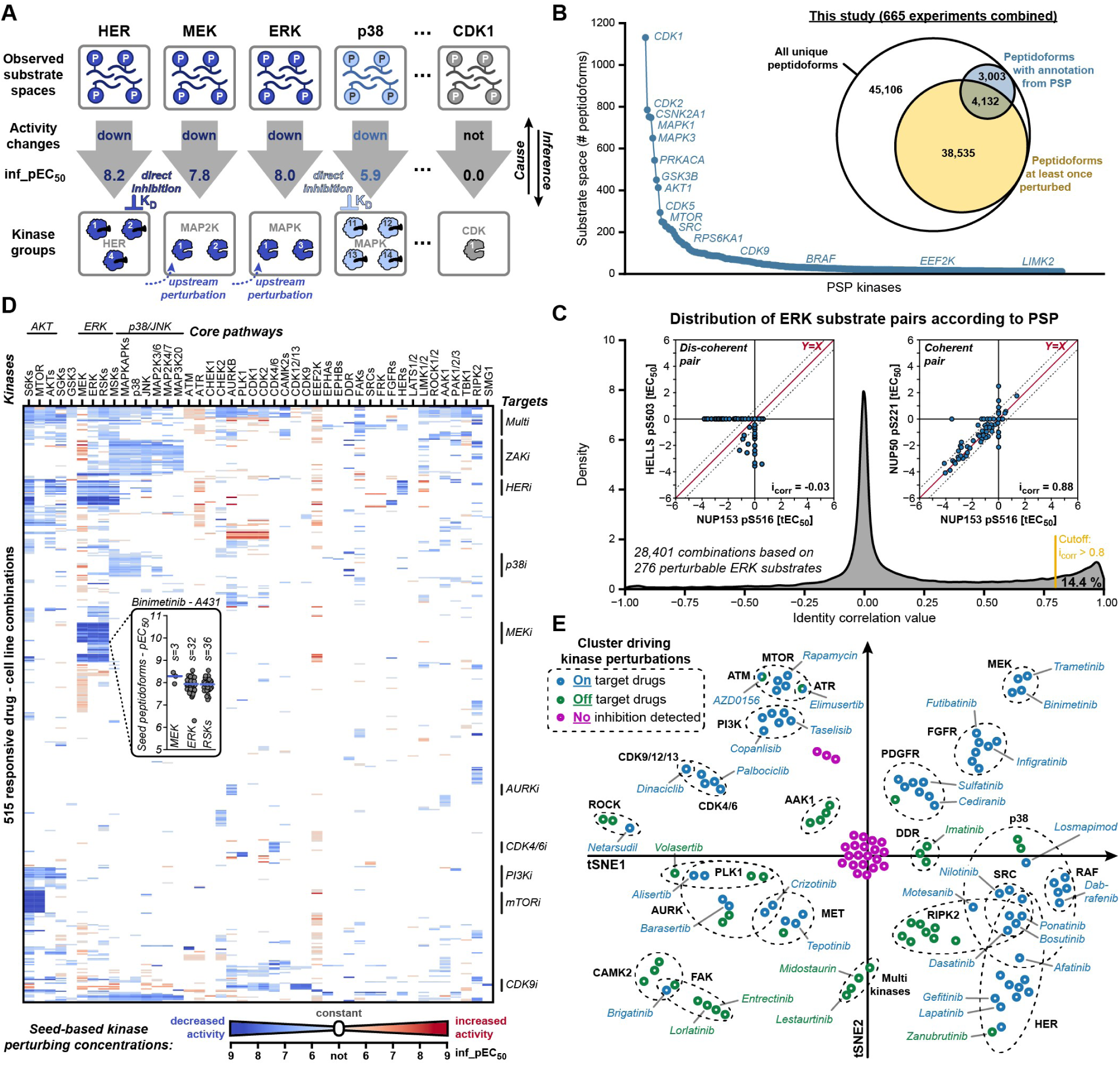
Kinase perturbation inference from its substrate space. ***(A)*** *Schematic representation of inferring the potency of kinase activity changes (expressed as an inferred drug concentration, inf_pEC_50_) in any one experiment from the median pEC_50_ of all of its observed regulated substrates in that experiment. Kinases are grouped if indistinguishable by known substrates. **(B)** Rank plot of the number of peptidoforms annotated as substrates of kinases according to PhosphoSitePlus (PSP). The inset Venn diagram compares the number of peptidoforms measured/regulated in this study with those for which kinase::substrate annotations are available in PSP. **(C)** Distribution of identity correlations for all pairwise ERK-substrate peptidoform combinations based on PSP annotation. Examples for coherent or dis-coherent peptidoform pairs are shown in the inset. The identity line X=Y and expected potency variation boundaries are indicated by red and black dotted lines, respectively. **(D)** Heatmap of inferred activity changes based on manually curated “seed” peptidoforms for 44 kinase groups across all decryptM experiments. The inset shows the pEC_50_ distribution of seed peptidoform for three kinase groups (s denotes the number of regulated seed peptidoforms) for A431 treated with Binimetinib as an example. **(E)** t-SNE projection of the heatmap, shown in panel D, grouped drugs (dots) by target class (dashed circles). Drugs are colored by whether or not cluster membership was due to on-target or off- target signaling*.

We next asked whether PSP-annotated substrate peptidoforms reliably monitor kinase activity changes in the potency dimension (*5*). If all annotations were correct, all substrates of the same kinase should have identical pEC_50_ within an experiment, and this potency coherence should exist across all experiments. To quantify the degree of potency coherence between a pair of peptidoforms across experiments (min. 4, max. 665), we developed the identity correlation value (i_corr_). First, pEC_50_ were transformed into tEC_50_ to distinguish peptidoforms that were up-, down-, or not-regulated in response to drugs on a harmonized scale with the highest drug dose set to zero. Thus, a tEC_50_ indicates relative potency and direction of change (positive for up, negative for down, zero for not). When all tEC_50_ pairs of two peptidoforms align along the identity line (Y=X), the peptidoforms are perfectly potency coherent (i_corr_=+1.0). If they align on the anti-identity line (Y=-X), they are perfectly anti-coherent (i_corr_=-1.0). If tEC_50_ mutually disagree and populate only the x- or y-axes, the peptidoforms are potency dis-coherent (i_corr_=0.0). To validate the i_corr_ approach, we analyzed all monophosphorylated peptidoform pairs that do vs. do not contain a missed tryptic cleavage site. These missed cleavages occur in any proteomic experiment because of incomplete tryptic digestion of proteoforms, share the same phosphorylation site, and thus represent ground-truth combinations. Among 2,040 such pairs, >80% showed strong potency coherence (i_corr_>0.8), whereas random combinations of peptidoforms (k=10,000) were tightly centered around zero, demonstrating expected dis-coherence (fig. S4AB).

Applying the potency coherence analysis to PSP-annotated substrate spaces of kinases revealed a substantial proportion of dis-coherence for many kinases. For example, only 14% of ERK substrate pairs had high i_corr_ values (fig. 3C). CDK1, AKT, MTOR, and p38 substrate pairs scored 7%, 14%, 24%, and 9%, respectively (fig. S4C- F). From this, we estimate that only 38% of ERK, 27% of CDK1, 38% of AKT, 49% of MTOR, and 30% of p38 annotations in PSP are potency-consistent and likely represent an upper bound of true KSRs. We suspect that the low potency reliability of PSP originated from the accumulation of many independent studies, such that incorrect substrate annotations overtook correct ones, resulting in a high global false discovery rate (FDR) and highlighting the need for appropriate KSR statistics (*15*).

### Curated seed peptidoforms enable estimating activity changes for 96 kinases

To overcome PSP’s low potency consistency, we manually curated 836 dynamic, kinase activity-indicating, and potency-coherent “seed” peptidoforms for 44 kinase groups, covering 96 kinases. For instance, MAPK1&3 were combined into an ERK group with 42 seed peptidoforms, and MAP2K1&2 into a MEK group containing MAPK1&3 activation loop phosphorylation. Seed pEC_50_ values for each kinase group were averaged to yield a single inf_pEC_50_ for each kinase group and drug experiment (fig. 3A). In Afatinib-treated A431 cells, we obtained inf_pEC_50_ for HER=8.2, MEK=7.8, ERK=8.0, RSKs=8.0, p38=5.9, and MAPKAPs=6.1 (fig. 3A, fig. S5). CDK1, CDK9, and GSK3 showed no activity change as most of their seed peptidoforms were not-regulated (fig. S5).

Calculating inf_pEC_50_ using seed substrates for all 44 kinase groups across all 665 experiments allowed us to systematically evaluate at which drug concentrations which kinases changed activity (fig. 3D). Several key findings emerged. First, kinase perturbation was relatively rare: 9% of all inf_pEC_50_ showed clear regulation, 74% showed absence of regulation, and 17% were inconclusive due to missing or unclear dose-response curves (fig. S6AB). Second, most inf_pEC_50_ were modest in potency (0.5-10 μM). As clinical inhibitors are generally potent for on-target kinases, these low-potency cases reflect off-target inhibition, consistent with in vitro affinity data (fig. S6CD)(*30*). This shows that off-target effects occur frequently in cellular perturbation assays when using kinase inhibitors at concentrations >500 nM and again highlights the importance of considering the potency dimension to distinguish inhibition events caused by different kinase targets. Third, downstream kinases were co-perturbed with always similar potencies (horizontal color shades in fig. 3D), clearly illustrated by the MEK-ERK-RSKs, MAP2K4/7-p38-MAPKAPK, AKTs-MTOR-S6Ks, and MAP2K3/6-JNK axes. Fourth, few kinases were frequently perturbed across experiments. Instead, most kinases were perturbed only in specific experiments (“islands” in fig. 3D), consistent with reports that inhibitors are often more selective in cells than in cell-free assays (*41*). This observation also reflects that not all kinases are active in all cell lines and, therefore, cannot be perturbed even if a particular inhibitor can, in principle, inhibit them. Fifth, the data revealed cell-line-specific pathway hierarchies. While upstream kinase inhibition led to potency-coherent downstream perturbation, downstream inhibition left upstream kinases unaffected (fig. S6E). Which upstream kinase was connected to which downstream kinase, e.g. ERK or AKTs, was different between cell lines (fig. S6F).

A t-SNE projection of the kinase potency matrix grouped drugs by target binding and pathway engagement, supporting the use of seed peptidoforms for inferring kinase activity changes (fig. 3E). Inhibitors of MEK, MTOR, PI3K, CDK, and p38 clustered tightly, as did inhibitors of RTKs such as HER, FGFR, PDGFR, and MET, because these kinases were active in at least one cell line. Some inhibitors clustered centrally, corresponding to weak responses or signaling not covered by the 44 kinase groups. Additional clusters emerged where on-target activity was low or absent and clustering was driven by strong off-target signaling. For example, the DNA-damage repair kinases ATM and ATR, despite being ubiquitously expressed (*42*), are typically weakly active in culture. Because ATM and ATR are evolutionarily related to MTOR and PI3K, the inhibitors AZD0156 (ATMi) and Elimusertib (ATRi) also target these kinases potently (*22*), explaining their clustering near the MTOR group.

### Potency coherence analysis identifies kinase::substrate relationships

We next applied the potency coherence concept to systematically mine decryptM data for substrates of the 44 kinase groups. The reasoning is as above: if a peptidoform is a unique and dynamic substrate of a kinase, its pEC_50_ must align with the inf_pEC_50_ of the phosphorylating kinase across all experiments (max. 665). This applies regardless of whether the kinase was directly inhibited as on- or off-target or was indirectly perturbed by inhibiting an upstream kinase. What matters is potency coherence between a kinase and its substrate across all experiments, quantified by i_corr_. We also developed a statistical approach to compute accurate p-values for each KSR hypothesis based on i_corr_ and the number of observations. Figure 4A illustrates this relationship between the kinase group S6Ks and its triply phosphorylated substrate RLS(ph)S(ph)LRAS(ph)TSK of the S6 ribosomal protein. Each point represents a dose-response experiment: the x-axis shows inf_tEC_50_ of S6Ks (derived from seed substrates), and the y-axis the observed tEC_50_ of the peptidoform. Strong potency coherence was observed (i_corr_=0.929, p-value=1.4e-81) across 457 dose-response curves. Extending this analysis to 19,084 peptidoforms that were responsive in ≥4 experiments revealed 209 peptidoforms with significant potency coherence (i_corr_>0.8, adjusted p<0.05), identifying them as S6Ks substrate candidates (fig. 4B). Across all 44 kinase groups (total: 839,696 KSR hypotheses), we identified 12,430 potency-coherent, candidate KSRs in this way (fig. S7AB).

**Fig. 4.**
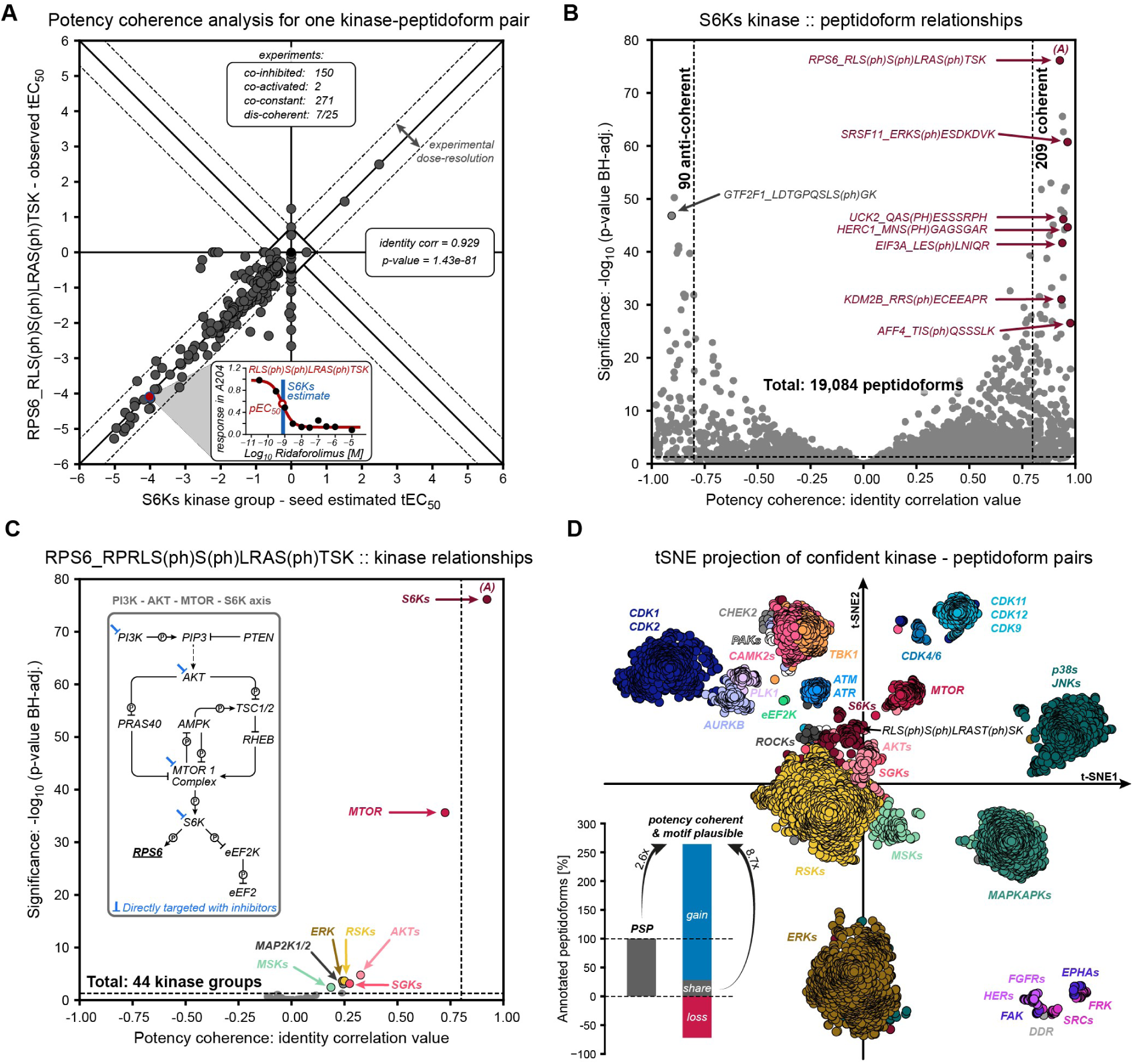
Annotating the human phosphoproteome with potency-coherent and motif-plausible kinase::substrate relationships. ***(A)*** *Potency coherence map akin to* fig. 3C*, but now correlating the activity change of a kinase group (here S6Ks) and one peptidoform (here RPS6_RLS(ph)S(ph)LRAS(ph)TSK) across all experiments (black dots). The inset shows the dose-response curve for this peptidoform in the Ridaforolimus-A204 experiment as an example (red dot in main panel). The vertical blue line indicates the estimated change in S6Ks activity from its seed peptidoforms in this experiment. (B) Volcano plot summarizing all 19,084 potency coherence maps for S6Ks::peptidoform combinations and highlighting several examples, including the map shown in A. Vertical dashed lines mark the chosen threshold of i_corr_>0.8, and the horizontal line marks the p- value_adj._<0.05 threshold. **(C)** Volcano plot akin to B, but this time asking which of the 44 kinase groups best explains the potencies of a peptidoform. The inset shows the canonical PI3K/AKTs/MTOR/S6Ks pathway. The pathway hierarchy is reflected in the volcano plot through decreasing order of statistical significance, as several upstream inhibitors of S6Ks were used. **(D)** t-SNE projection of all confident (potency-coherence and motif-plausible) kinase::substrate relationships colored by kinase groups. The inset quantifies the gains and losses in kinase::substrate relationships relative to PSP resulting from the data analysis presented in this work*.

Visualizing the coherence analysis as a volcano plot for one kinase against all perturbed peptidoforms showed the overall impact of that kinase on the dynamic phosphoproteome. For example, RIPK2 (19 KSRs) and ATM (61 KSRs) had few coherent peptidoforms, whereas MTOR (183 KSRs) and RSKs (1,497 KSRs) exhibited a much larger impact (fig. S8). From the substrate perspective, volcano plots also show how well each kinase group explains the tEC_50_ pattern of a specific peptidoform. For RLS(ph)S(ph)LRAS(ph)TSK, only S6Ks scored above the i_corr_ and p-value thresholds, while upstream kinases MTOR and AKTs did not (fig. 4C). All potency coherence maps and volcano plots (fig. 4A-C) were integrated into ProteomicsDB (*23*) (proteomicsdb.org/analytics/KSR), enabling interactive exploration of KSRs at the kinase-, protein-, or peptidoform-level. These tools were implemented such that future decryptM experiments can be automatically integrated, and each KSR will be re- evaluated based on the new collective evidence. This marks an important improvement in KSR mining because, for the first time, there is a clear data- and statistics-driven mechanism to update KSR annotations while controlling global FDR.

### Potency coherence and motif plausibility are orthogonal criteria to constrain kinase::substrate relationships

For many peptidoforms, potency coherence analysis identified one significant kinase for one peptidoform (1::1 KSRs, fig. S7C). This was possible because the used drugs included inhibitors acting at different positions in linear signaling pathways, generating distinct perturbation profiles and enabling, for example, separation of kinases within the PI3K-AKTs-MTOR axis (fig. 4C). However, some kinases were never directly targeted by any drug but perturbed only indirectly via signaling cascades. This produced similar potency values for kinases along the same pathway, making them indistinguishable in the potency dimension. Therefore, ERK vs. RSKs and p38s vs. MAPKAPKs KSRs were difficult to separate by potency alone (n::1 KSRs, fig. S7C). This is not a general limitation of potency coherence analysis, but reflects the restricted target space of clinical kinase inhibitors used here. Extending decryptM profiling to chemical probes for further kinases will overcome this issue. For now, we leveraged the property that many linear signaling cascades involve kinases from different families (*3*). ERK and p38, for instance, are CMGC-family kinases with proline-directed motifs, whereas RSKs (AGC-family) and MAPKAPKs (CaMK-family) prefer basophilic motifs. Thus, kinase motifs provide an orthogonal criterion to exclude potency-coherent but motif-implausible assignments. Incorporating data from the Kinase Library (*18, 19*) successfully separated ERK from RSKs and p38 from MAPKAPKs (fig. S9AB). The reverse was not always true, i.e. many kinases that could be clearly separated by potency could not be distinguished by motif analysis alone (fig. S9CD).

Systematically combining potency coherence and motif plausibility improved the separation of candidate KSRs (projected as 2D t-SNE; fig. 4D) and yielded 5,318 “confident” (i_corr_>0.8, adjusted p<0.05, motif score>0.0) KSRs across 44 kinase groups and 1,809 proteins. This represents a ∼2.6-fold gain over PSP-based annotations for the same kinases. Notably, only ∼31% of PSP annotations remained when requiring both potency coherence and motif plausibility, consistent with our initial assessment above. Prominent examples of false PSP assignments are AKTs substrates that are, in fact, S6Ks or RSKs substrates. Overall, our analysis increased confident KSRs for 44 kinase groups by ∼8.7-fold. These re-annotations should immediately enhance kinase activity inference tools that rely on accurate KSRs.

### Coherence analysis identifies signal executors, transducers, and integrators

We observed that kinases differed markedly in substrate numbers (fig. S10A). Eight kinase groups phosphorylated ten or fewer proteins and may be viewed as local “executors” with specific cellular tasks. Examples include EEF2K phosphorylating EEF2, a regulator of protein synthesis; LATS1/2 phosphorylating AMOT, a central Hippo pathway component; and CDK4/6 phosphorylating RB1, RBL1, and RBL2, which control cell-cycle progression from G1 to S phase. In contrast, 11 kinase groups targeted over 100 proteins, acting as major “signal transducers.” Among them were RSKs, ERK, and JNK (410, 498, and 273 substrate proteins, respectively), which are known to shuttle between cellular compartments to deliver their signals throughout the cell (*43*).

We next analyzed how many kinases phosphorylate a given protein and observed an exponential decline, indicating that most proteins are dynamically regulated by only one or two kinases (fig. S10B). Still, 104 proteins appeared to function as signal integrators because they were phosphorylated by five or more kinases at distinct sites. Classic integrators included TSC2 (6 kinases) and IRS1 (9 kinases) of the MTOR pathway (fig. S10C) (*44, 45*). Additional examples were MDC1 (9 kinases), a DNA damage repair scaffold, and MKI67 (8 kinases), a clinical tumor marker. Novel integrators included large cytoskeleton–membrane linkers such as AHNAK (686 kDa) and PLEC (517 kDa), phosphorylated by 12 and 8 kinases, respectively, and consistent with the role of kinases in cytoskeletal remodeling (fig. S10D).

### Potency coherence connects genotypes, proteotypes, and phenotypes of cancer cells

Predicting phenotypic outcome of kinase inhibition from genomic data has been difficult because cancer signaling is highly cell-context-specific, shaped by combinations of genetic alterations, differentiation states, protein expression, PTM levels, and proteoform turnover, to name a few. We reasoned that the potency dimension of decryptM and confident KSRs could help bridge this genotype-phenotype gap, because potency is a characteristic inherent to a drug-target pair, irrespective of the assay type. We therefore integrated the genomic data from the DepMap program (*31*) and dose-dependent viability data for 144 inhibitors, recently published by the authors (*46*), with the decryptM profiles acquired here (fig. 5A; fig. S11). This analysis provided three important pieces of information: First, the decryptM profiles revealed cell-line-specific kinase signaling pathways (“connectivity” is indicated as red arrows in fig. 5A); Second, they measured the overall impact of a specific kinase on the phosphoproteome (“perturbation extent” is indicated as red bars in fig. 5A). Third, they molecularly rationalized observed viability changes by associating kinase and peptidoform potencies with phenotype potencies (fig. S11).

**Fig. 5.**
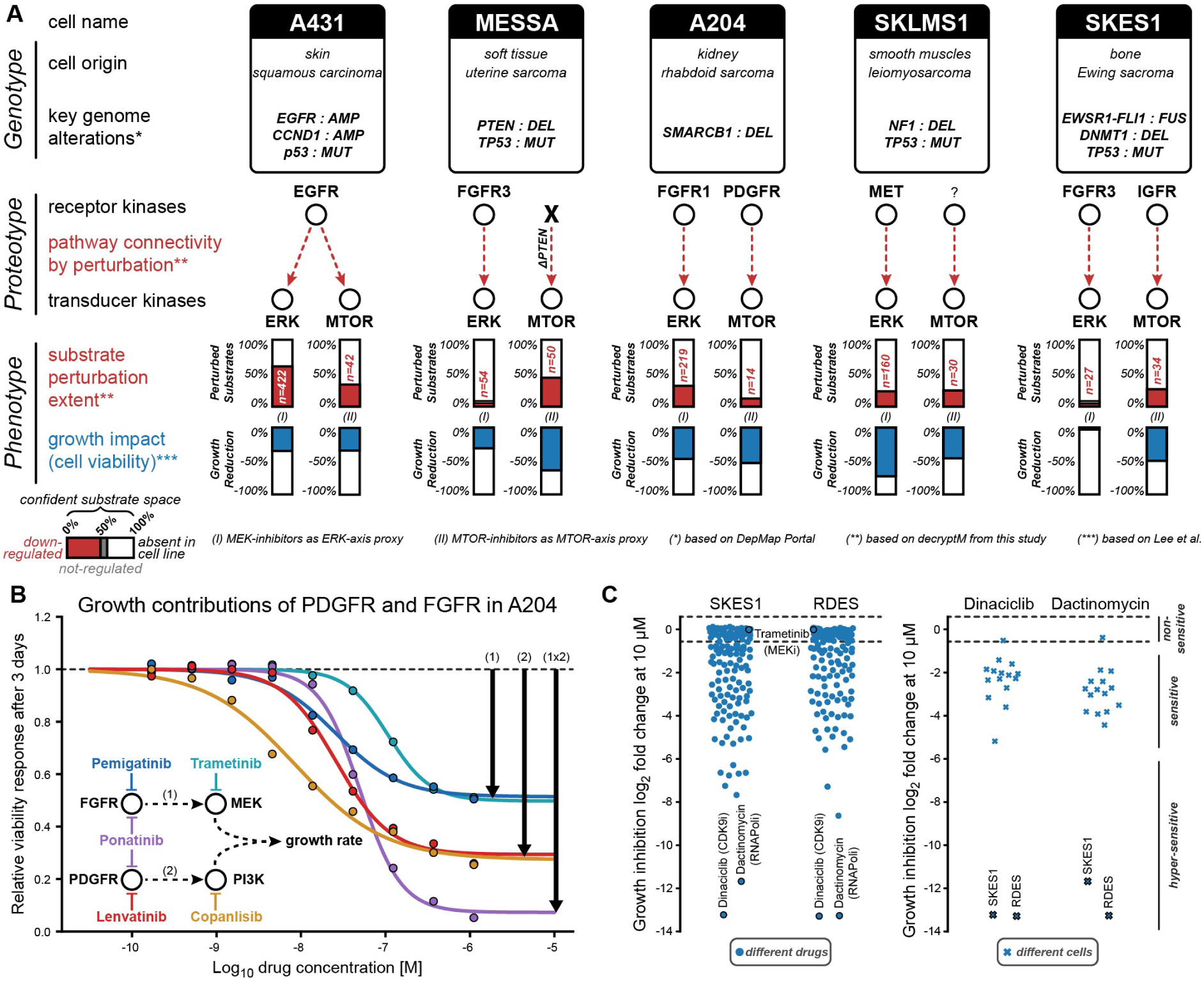
Potency coherence connects genotypes to proteotypes to phenotypes. ***(A)*** *Schematic representation of relating genotype (alteration data from DepMap Portal: AMP=amplification, MUT=mutation, DEL=deletion, and FUS=fusion), proteotype (decryptM: data from this study), and phenotype (viability data from Lee et al.) for five cell lines and using the ERK and MTOR axes as examples. Receptor to transducer kinase connectivity (red dotted arrow) was determined by potency-coherent perturbation patterns of the most upstream kinase (e.g. see CurveCurator dashboards for EGFR:Afatinib, FGFR:Futibatinib, PDGFR:Lenvatinib, MET:Tepotinib, IGFR:Linsitinib). The extent of kinase activity perturbation in each cell line (colored bars) is depicted by the number (and percentage of total) of confident ERK and MTOR substrates that were down-regulated (red), not-regulated (gray, barely visible), or absent/unclear (white). For phenotypic (cell viability) measurements, Trametinib was used as an ERK-axis proxy and Rapamycin as an MTOR-axis proxy. **(B)** Cell viability dose-response curves for Pemigatinib, Trametinib, Ponatinib, Lenvatinib, and Copanlisib in A204. Black bold arrows at the back plateaus indicate the contribution of each axis alone* (*1*) *&* (*2*) *or combined (1x2) to cell growth inhibition. The simplified pathway in the inset depicts the inhibitors, their targets, and kinase connectivity in A204. **(C)** Swarm plots showing growth inhibition of SKES1 and RDES in response to 144 inhibitors, notably insensitivity to Trametinib (MEKi; left panel) but hypersensitivity to Dinaciclib (CDK9i) and Dactinomycin (RNAPoli; right panel)*.

To illustrate the above, we focused on ERK and MTOR signaling and found that they often acted independently and were regulated by different RTKs, except in A431, where both ERK and MTOR were regulated by amplified and overactive EGFR. In this cell line, the high EGFR activity translated into high ERK and MTOR activity, reflected by many down-regulated, potency-coherent, and motif-plausible substrates (perturbation extent: 422 of 715 ERK substrates; 42 of 127 MTOR substrates; all identified in this work). We note that the extent of drug-regulated substrates drastically varied between cell lines, indicating different levels of kinase activity in these cells (fig. 5A). In the sarcoma cell lines A204 (rhabdoid), SKES1 (Ewing), and MESSA (uterine), ERK was connected to FGFR. In SKLMS1 (leiomyosarcoma), ERK was regulated by MET. MTOR was PDGFR-dependent in A204 and IGFR-dependent in SKES1. In MESSA, however, a PTEN loss-of-function mutation (*47*) resulted not only in pronounced MTOR activation (perturbation extent of 50 substrates), but also in decoupling from upstream RTKs as no RTK inhibitor decreased AKTs or MTOR activity (Fig. S12A) or cell growth at relevant potencies (Fig. S12B). Because the PI3K inhibitor Copanlisib strongly perturbed AKTs and MTOR, the decryptM profiles clearly verified decoupling at this position and explained the growth phenotype.

While EGFR overactivity in A431 and AKTs/MTOR-decoupling in MESSA were, in principle, genetically predictable, most sarcomas exhibit low mutational burden, few amplifications, deletions, or fusions (*48*). This makes deducing kinase activity dependencies for therapy recommendations non-trivial. For example, a genomic deletion of the tumor suppressor SMARCB1 disrupts the chromatin remodeling complex SWI/SNF in A204 (*47, 49*), resulting in a genomically unpredictable but phenotypically observed dependence on both PDGFR and FGFR (upstream of MTOR and ERK respectively according to decryptM data; fig. 5A). As a consequence, single-axis inhibition only induced partial growth reduction but applying Ponatinib, blocking both RTKs, yielded a multiplicative response (fig. 5B). A similar multiplicative response characteristic for inhibiting the ERK and MTOR axes was observed in A431 (fig. S12C).

Another example for which genomic information cannot predict kinase activities or connectivity is the cell line SKES1, which carries the EWSR1-FLI1 transcription factor fusion (*47, 50*). DecryptM profiling showed minimal FGFR-ERK signaling (perturbation extent: 27 of 715 ERK substrates), explaining the absence of phenotypic responses to ERK-axis inhibition. Instead, only IGFR inhibition suppressed growth via MTOR (fig. 5A). The same phenotypic patterns were observed in RD-ES, another EWSR1–FLI1- positive cell line (*47*). In both, the strongest phenotypic responses were, however, observed for transcriptional inhibitors, such as Dactinomycin or the CDK9 inhibitor Dinaciclib, which act downstream of the fusion and clearly distinguished them from other lines (fig. 5C). These results indicate a high level of transcriptional addiction to the fusion protein, and rationalize the observed insensitivity to ERK inhibition and CDK9 vulnerability. Ongoing clinical trials will show whether this vulnerability also translates into a clinical benefit (*51*).

Overall, potency coherence provided a rational framework for aligning context-specific signaling with inhibitors targeting relevant cancer pathways. Conceptually, this could improve precision oncology by i) identifying effective drugs, ii) excluding ineffective ones, and iii) justifying the use of multi-kinase inhibitors despite potential toxicity. Yet, realizing this potential requires accurate deduction of patient signaling states from biopsy material.

### Inferring kinase activities with confident KSRs in cancer patient tumors

As part of ongoing pan-cancer precision oncology programs (*52–55*), we have measured the (phospho)proteomes of >2,000 advanced cancer patients. We hypothesized that confident KSRs identified in this study could be used to detect kinase activities in individual patients. After grouping patient-derived peptidoforms, batch normalization, and removal of protein expression contributions (fig. S13), we retained 16,933 peptidoforms with >67% data completeness for 1,627 patients with >30% tumor-cell content. Comparing dynamic peptidoforms (≥4-times regulated) in cell lines with those in biopsies revealed ∼27% overlap (fig. 6A). Of these, 1,204 (16%) met our confident KSR criteria. We then calculated kinase activity scores by aggregating substrate abundances into kinase z-scores (fig. 6B; fig. S14A). Positive z-scores indicate that most substrates had elevated abundance, near-zero values indicate average abundance, and negative values indicate reduced abundance. For example, patient P1038 showed very high ERK and RSKs activity, while P2485 exhibited high CDK2 activity. Correlating z-scores across the cohort confirmed their biological relevance: kinases within the same pathway (e.g., ERK/RSKs, p38/MSK) correlated well (fig. 6C; fig. S14B), whereas kinases from unrelated pathways, such as ATR vs. ERK, did expectedly not. Kinases that are co-activated during specific cell- cycle phases, such as CDK2 and AURKB, also showed strong agreement. We conclude that confident KSRs from cancer models can be used for capturing kinase activity states in patient biopsies, where only a single abundance measurement is available per patient.

**Fig. 6.**
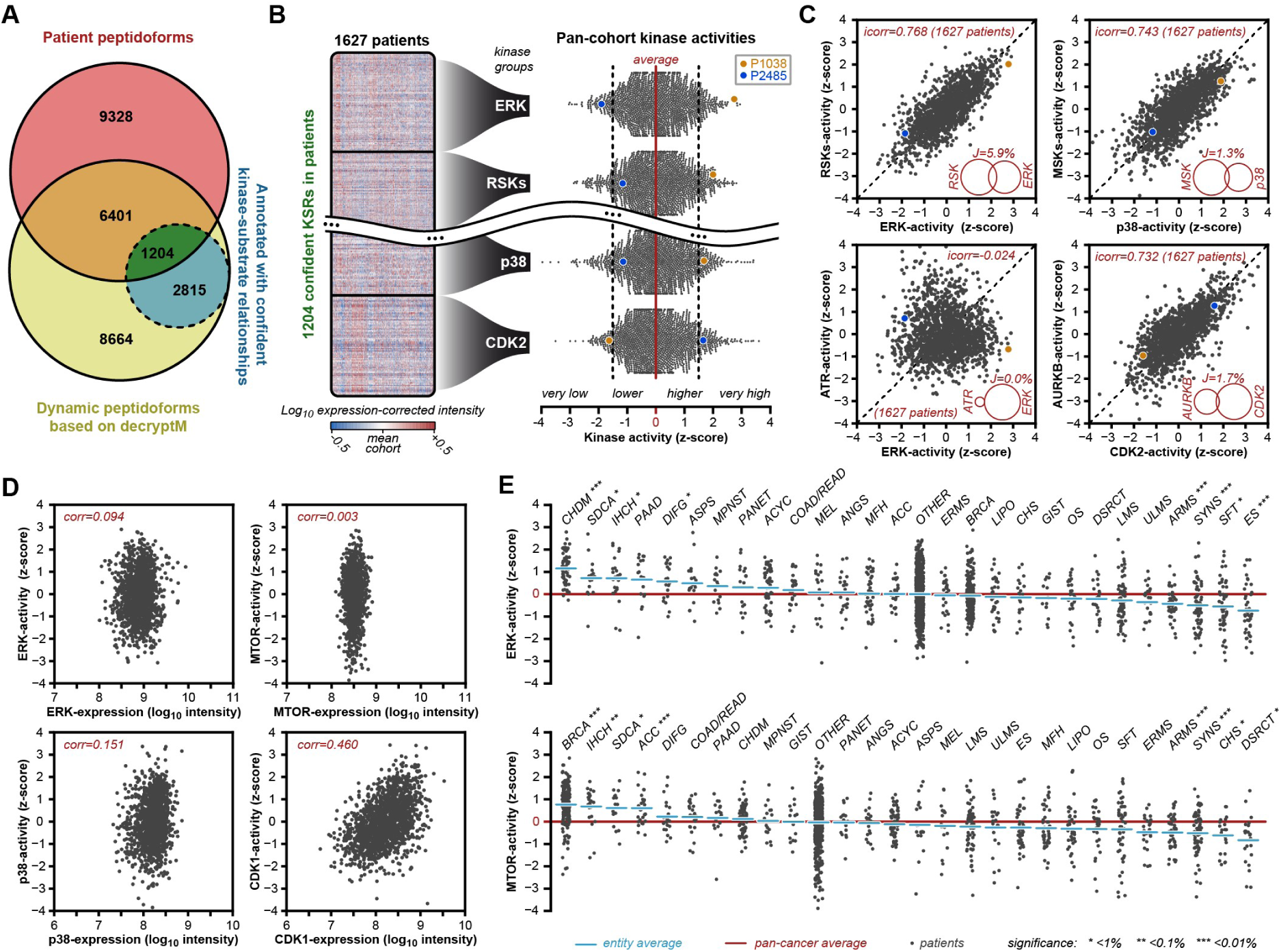
Kinase activity scoring of cancer patient biopsies. **(A)** Venn diagram showing the overlap between phosphorylated peptidoforms identified in the clinical cohort, dynamic peptidoforms by decryptM, and confident KSRs. **(B)** Examples of kinase activity analysis in patient samples by aggregating the abundance of KSRs (intensity heatmap) into a single activity z-score for each kinase group (here ERK, RSKs, p38, CDK2) and patient. The swarm plots show the distribution of kinase activities of all patients in the cohort. Two patients, P1038 (orange) and P2485 (blue), are highlighted as examples. **(C)** Identity correlation analysis of two kinase activities (z-scores), highlighting the same two patients as in B. Venn diagrams in the insets show the overlap between substrates of the compared kinase groups. **(D)** Pearson correlation analysis of kinase expressions and activity across patient samples for ERK, MTOR, p38, and CDK1. **(E)** Swarm plots of ERK (upper panel) and MTOR (lower panel) activity in patient samples grouped by cancer entity. The average pan-cancer activity is indicated as a red line. The entity-specific average activity is shown as blue lines. Entities were sorted by average activity. Significantly increased or decreased activity was determined using a two-sided one-sample t-test. Statistical confidence is indicated as * (p<1%), ** (p<0.1%), and *** (p<0.01%). The entity abbreviations follow the OncoTree nomenclature. Entities with N<15 patients were grouped as OTHER.

We next asked whether kinase activity could be inferred from protein expression alone. Using ERK, MTOR, p38, and CDK1 as examples, activity scores showed little or no correlation with protein expression (fig. 6D). This challenges a common assumption that high kinase expression implies high activity (*17, 56*). While overexpression can activate some RTKs that dimerize at pathological concentrations, most intracellular kinases require elevated upstream signaling for activation. This provides a clear rationale for systematically including phosphoproteomics in the analysis of clinical samples.

Finally, we applied confident KSRs to compare kinase activity between cancer entities. Chordomas (CHDM) displayed significantly higher ERK activity than the pan-cancer cohort, while several carcinomas (BRCA, IHCH, SDCA, ACC) showed higher MTOR activity (fig. 6E). In contrast, sarcomas that are defined by transcription-factor fusions (e.g., EWSR1-FLI1 in ES, NAB2-STAT6 in SFT, SS18-SSX in SYNS, PAX3-FOXO1 in ARMS, (*57*)) showed significantly lower ERK activity, consistent with our cell-line observations above. This suggests that ERK-axis decoupling occurs similarly in patients with transcription-factor-fused sarcomas. Still, individual patient kinase activities varied substantially within an entity, possibly reflecting additional genomic (or other) alterations acquired during therapy that increased kinase activity in these advanced patients. We anticipate that the kinase activity estimation as presented here will become a valuable resource for molecular tumor boards, clinical trials, and eventually routine diagnostics. However, treatment success will depend on a systematic understanding of kinase activity in each cancer context. The potency- coherence concept thus provides a strong theoretical and practical foundation for adding more precision to precision medicine in the future.

## Discussion

We introduced a new concept, termed potency coherence analysis, for identifying KSRs. Its successful implementation was based on the systematic, dose-dependent perturbation of cellular kinase activities using 133 well-characterized clinical kinase inhibitors, resulting in 17.1 million peptidoform dose-response profiles measured by mass spectrometry. The core idea is that the dose-response profile of any phosphorylation site reflects a change in kinase activity and that the two are causally linked in the potency rather than the effect size dimension. Because many kinase inhibitors target multiple kinases with varying potencies, our statistical approach discovered 5,318 confident KSRs for 96 human kinases. A particular strength of the approach is that all interpretable dose-response peptidoform profiles, whether up-, down-, or not-regulated, were used. The value of negative data is often overlooked in perturbation studies, but represents a distinct advantage of high-throughput technologies over classical biochemical approaches. The potency-coherent and motif- plausible KSRs established here have already proven useful for understanding signaling in cell lines, explaining growth phenotypes in response to drugs, identifying errors in public databases, and estimating kinase activities in patients with diverse cancers. The latter is especially promising in precision oncology and drug discovery, where these confident KSRs could serve as pharmacodynamic biomarkers that directly link a drug’s mode of action to its target activity in patients.

This study represents substantial progress in establishing a statistical and analytical framework for perturbation-based high-throughput analysis of KSRs. Future collaborative efforts within the scientific community can build upon this foundation to address the remaining limitations of this study. Although we identified thousands of highly confident KSRs, most phosphorylation sites still lack rigorous annotation. Several factors contributed to this: first, we focused on clinical-grade kinase inhibitors for their translational potential in oncology. However, these drugs do not cover all 518 human kinases. Incorporating additional inhibitors and chemical probes developed by the chemical biology community (*58, 59*) will help close this gap. Second, our analysis was limited to five cell line models, which do not capture the full range of kinase activities and possible substrate proteins. This can be addressed by expanding decryptM to larger cancer cell line collections such as DepMap or GDSC (*31, 60*). Third, we focused on dynamic substrates using a 100-minute treatment window, which biased the analysis against phosphorylation events with slower kinetics and prevented linking kinases to static phosphorylation sites that may serve important structural roles (*61*). Fourth, the current literature provides no or few reliable seed sites for understudied kinases to infer perturbation potencies. While the first three limitations mainly require additional effort and capacity, overcoming the fourth will likely need tailored strategies such as engineering substrate-trapping mutants (*62*) and developing specialized cell models for kinases with narrow physiological substrate spaces.

Because potency is a universal physicochemical property of drug-target interactions, integration of future decryptM data into this framework will be seamless and may ultimately yield unique substrates for every human kinase. Although potency coherence and motif plausibility provide rigorous, orthogonal criteria, the overall success of the approach depends on progressively falsifying KSRs, ideally until only one confident kinase group remains for each peptidoform. This will systematically eliminate alternative (and currently unknown) kinase hypotheses, which are often ignored and have led to overinterpretation in the past, particularly for overstudied kinases. Thus, the 5,318 KSR annotations reported here should be regarded as the best current explanations, derived from 665 experiments with 133 inhibitors in five cell lines. Continued perturbation of diverse systems with well-characterized inhibitors will be essential for correcting and further refining our understanding of cellular signaling and kinase biology.

Realizing this vision also requires continuously updating, re-evaluating, and falsifying KSRs in public databases. To this end, we have implemented the potency coherence analysis in ProteomicsDB (*23, 63*), so that KSRs are re-evaluated whenever new decryptM datasets are uploaded. A true, unique, and dynamic substrate must change abundance at the potency at which its causal kinase has changed activity. False-positive annotations are increasingly penalized as they fail to explain potencies across experiments. True KSRs may also be penalized because our simplified assumptions do not cover the possibility of shared or conditional KSRs. This logic is akin to particle physics, where deviations from the “standard model” lead to refinements of the theory. We propose it is time for a “standard model of cellular signaling,” which may be built by moving away from low-throughput, hypothesis-driven studies toward systematic, high-throughput, PTM-centric, and fully quantitative approaches. The potency coherence analysis offers a unifying framework for continuously mapping kinase or more broadly any enzyme-substrate relationship while rigorously controlling global false discoveries at the same time.

## Funding

This work was partly funded by the Federal Ministry of Education and Research (BMBF: CLINSPECTM, DROP2AI–031L0305A, HEROES-AYA–01KD2207F), the German Research Foundation (DFG: SFB1309–325871075; SFB1335–360372040; PhoSAIC–537476536), the European Research Council (ERC: TOPAS–833710), and the European Union’s Horizon 2020 Program (grant agreement no. EPIC-XS– 823839).

## Author contributions

FPB, MT, and BK developed the concept of potency coherence analysis. FPB, NK, MA, YCC, and CYL performed laboratory experiments. FPB, JM, LDED, CJ, FH, AS, and MT performed data analysis and data integration. CYL, AS, MT, and BK directed and supervised experiments and data analysis. FPB and BK wrote, reviewed, and edited the manuscript with input from all authors.

## Competing interests

BK is a founder and shareholder of MSAID. He has no operational role in the company. All other authors declare no competing interests.

## Data and materials availability

All raw data, supplement data/text, and source code will be made available with the peer-reviewed journal publication. The potency coherence analysis is available as a new Python package, called “icorr”, on GitHub (github.com/kusterlab/icorr) and PyPi (icorr). Interactive data exploration is implemented in proteomicsDB for dose-response curves (proteomicsdb.org/decryptm) and KSRs (proteomicsdb.org/analytics/KSR).

---

**Figure S1:**
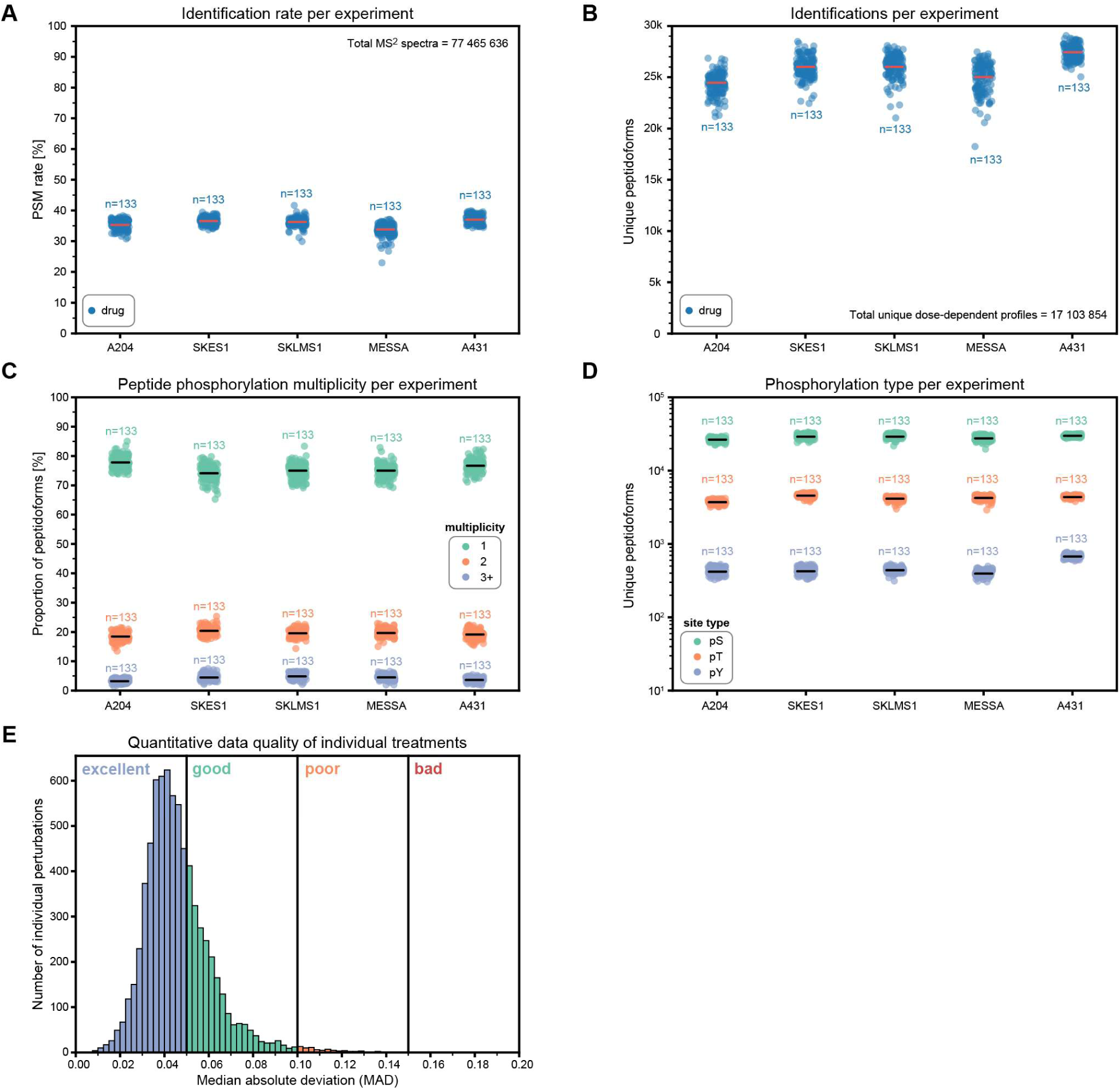
Different data quality metrics for all decryptM datasets. ***(A)*** *Identification rate of MS-spectra separated by cell lines. Each dot is an independent decryptM experiment. The mean PSM rate is indicated as a line. **(B)** Unique peptidoforms identified in each experiment, separated by cell lines. The mean peptidoform number is indicated as a line. **(C)** Multiplicity analysis per experiment (1 phosphorylation: green, 2 phosphorylations: orange, 3+ phosphorylations: blue) of the different peptidoforms separated by cell lines. The mean proportion is indicated as a line for each multiplicity group. **(D)** Site type analysis per experiment (phosphorylation on serine (pS): green, phosphorylation on threonine (pT): orange, phosphorylation on tyrosine (pY): blue) of the different peptidoforms separated by cell lines. The mean count is indicated as a line for each site type group. **(E)** The distribution of CurveCurator’s noise analysis of all 7,315 individual treatments is separated into 4 quality areas (excellent, good, poor, bad). The median absolute deviation (MAD) value indicates how far 50% of all observation ratios are with respect to the estimated curve model for a specific dose*.

**Figure S2:**
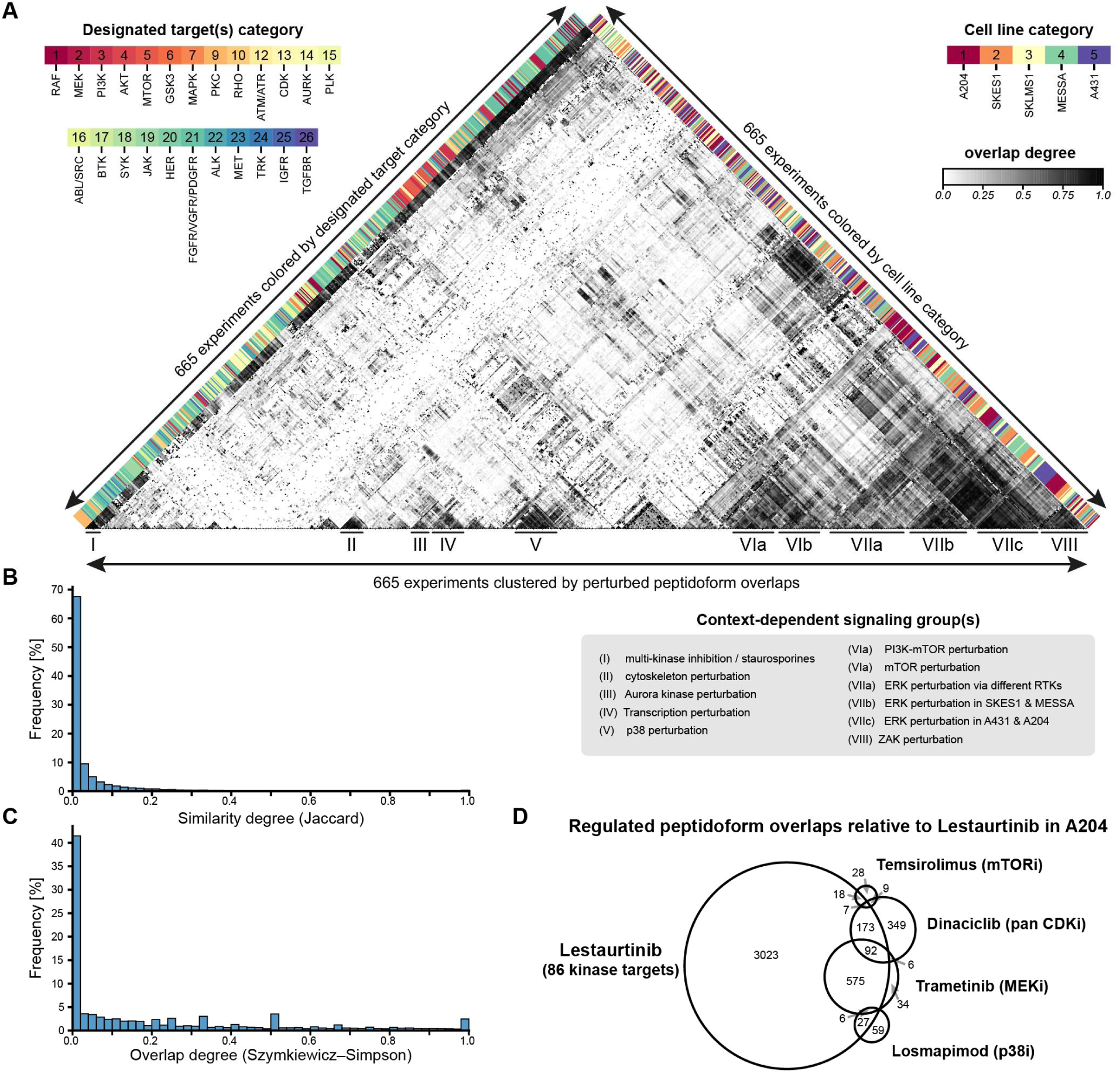
Comparisons of drug-regulated peptidoform sets. ***(A)*** *Pairwise overlap comparison of all 665 decryptM experiments identifies peptidoform perturbation signatures after clustering (horizontal bottom line). Each experiment is a specific combination of a drug and a cell line. Peptidoforms with unclear regulation classification were excluded from the analysis. The overlap degree between two experiments (E_i_, E_j_) was measured using the Szymkiewicz–Simpson index conditioned by the regulation classes up, down, and not. The resulting overlap value is gray-scaled in the heatmap. The cell line and drug categories were color-coded along the axis. The drug color code is based on conventional drug classes, which are primarily defined by the designated target rather than off-targets or all targets. Formed signaling clusters correspond to overlapping peptidoform signatures (enumerated from 1 to 8) along the horizontal line. Cell line or drug categories did not form global clusters. **(B)** The distribution of similarity degrees of all ∼221k combinations indicates very few redundant (high similarity) experiments. **(C)** The distribution of overlap degrees of all combinations indicates much more overlap in the datasets compared to the similarity of Fig. S2B. **(D)** Example of peptidoform containment (overlaps) in the sarcoma cell line A204 between Lestaurtinib, Dinaciclib, Trametinib, Losmapimod, and Temsirolimus perturbations*.

**Figure S3:**
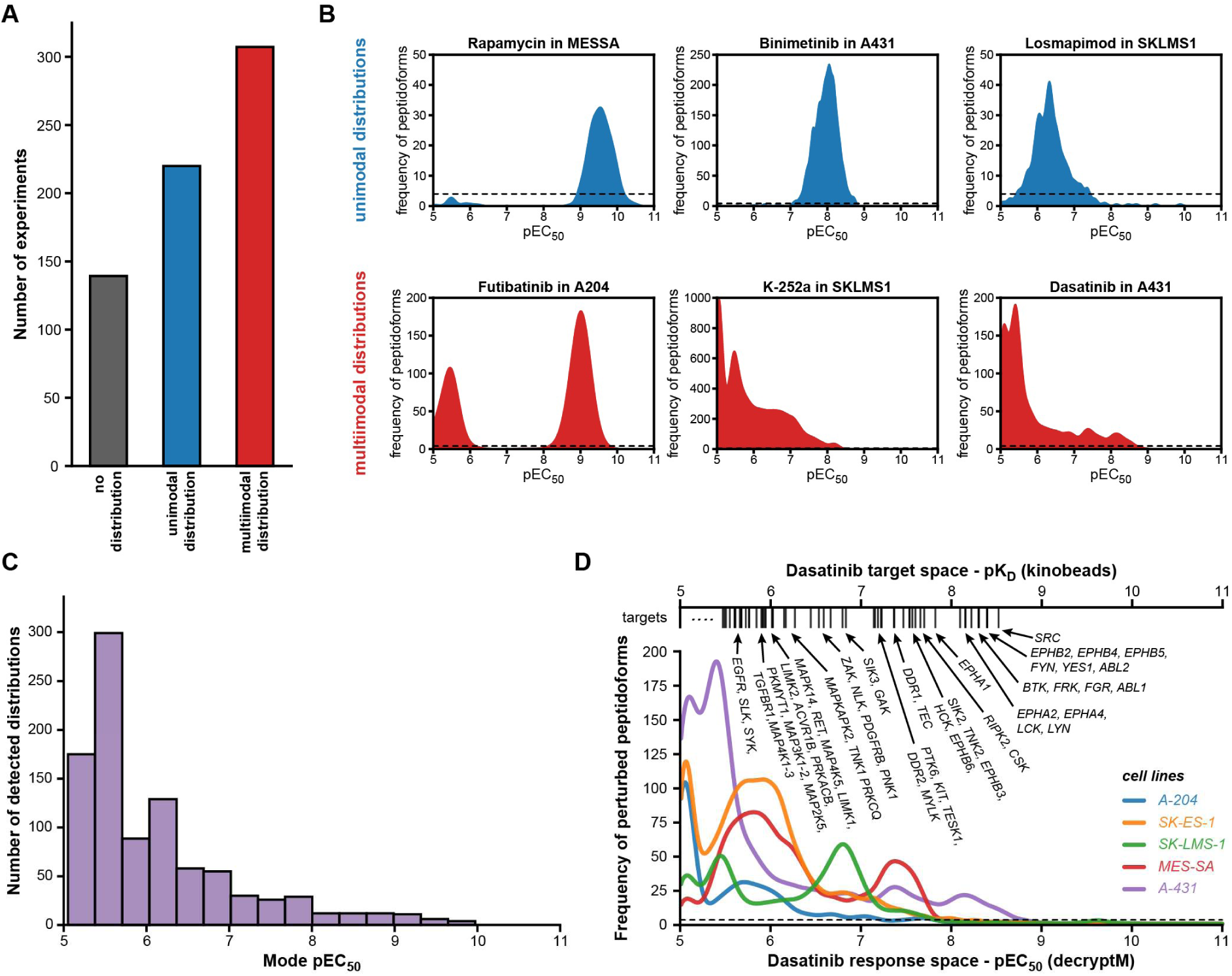
Polypharmacology is commonly observed in decryptM pEC_50_ peptidoform distributions. ***(A)*** *Bar chart to show how often uni- or multimodal distributions exist in the 665 decryptM experiments. Distribution detection was performed by first fitting a Gaussian KDE and then applying SciPy’s find_peaks algorithm. Experiments were then grouped and counted if they had no (gray), one (blue), or multiple (red) distinct peptidoform distributions in the potency dimension. Each distribution indicates at least one target engagement. **(B)** Six example distributions for unimodal (blue) or multimodal (red) distributions. The dashed line is the minimal required frequency to detect a peak. **(C)** Mode pEC_50_ distribution of all detected modes over all 665 experiments. Many responses exist in the off-target area 1-10 µM. **(D)** pEC_50_ distributions for a single drug, Dasatinib, in five different cell lines (colored). A second x-axis on top of the plot shows the pK_D_ values and corresponding kinases that Dasatinib can directly bind and inhibit, as was determined by Kinobeads (Klaeger et.al). Not only does Dasatinib have multimodal pEC_50_ distributions within all cell lines, but it also has different pEC_50_ distributions across cell lines, which are dependent on the present kinase activities in these cancer cell models. In short, the drug determines at which concentration a potential response could happen, while the specific cell biology determines if a response is possible*.

**Figure S4:**
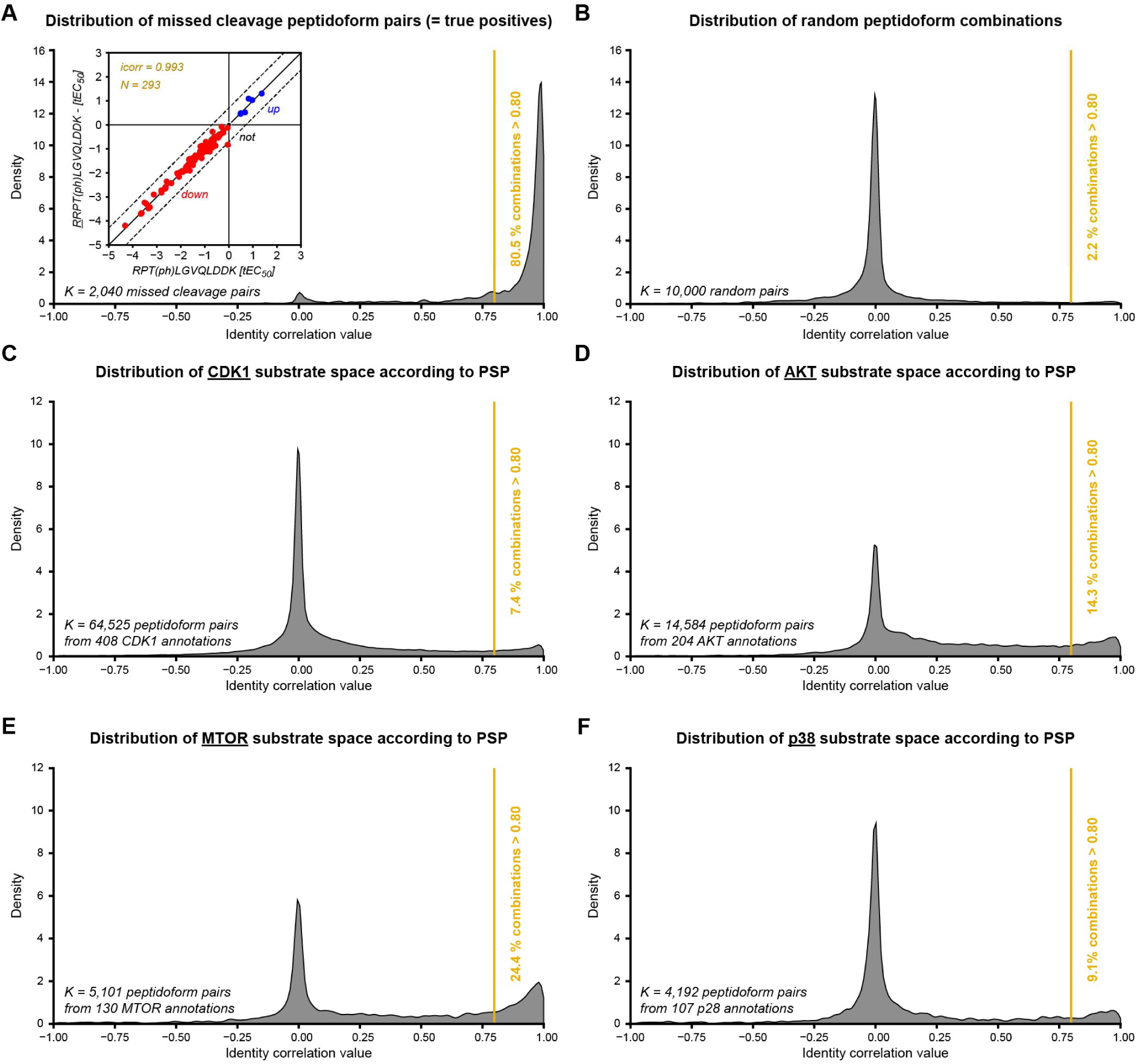
Consistency analysis of the potency dimension of substrate spaces of PhosphoSitePlus (PSP) kinase annotations. ***(A)*** *Identity correlation was analyzed for 2,040 missed-cleavage pairs over all experiments. Because missed-cleavage pairs capture the same sites, they must have the same pEC_50_ value within the error of dose resolution and measurement variance. The identity correlation analysis is exemplified with the missed-cleaved peptidoform pair RPT(ph)LGVQLDDK - RRPT(ph)LGVQLDDK from the THOC5 gene. Transformed EC_50_ (tEC_50_) values are defined as the pEC_50_ value relative to the experimental dose range. Down-regulation is indicated by a (-) sign, and up-regulation is indicated by a (+) sign. **(B)** Identity correlation was analyzed for 10,000 random peptidoform pairs from the entire dataset. Pairs from the same protein were excluded from the analysis due to potential shared dependencies. **(CDEF)** Identity correlation was analyzed for different kinase substrate spaces according to PSP. Pairs with fewer than three perturbations were excluded from the analysis. The identity correlation threshold was set to 0.8 (yellow line). To estimate the percentage of consistent annotations in the substrate space of a kinase, we took the square root of the coherent pair proportions*.

**Figure S5:**
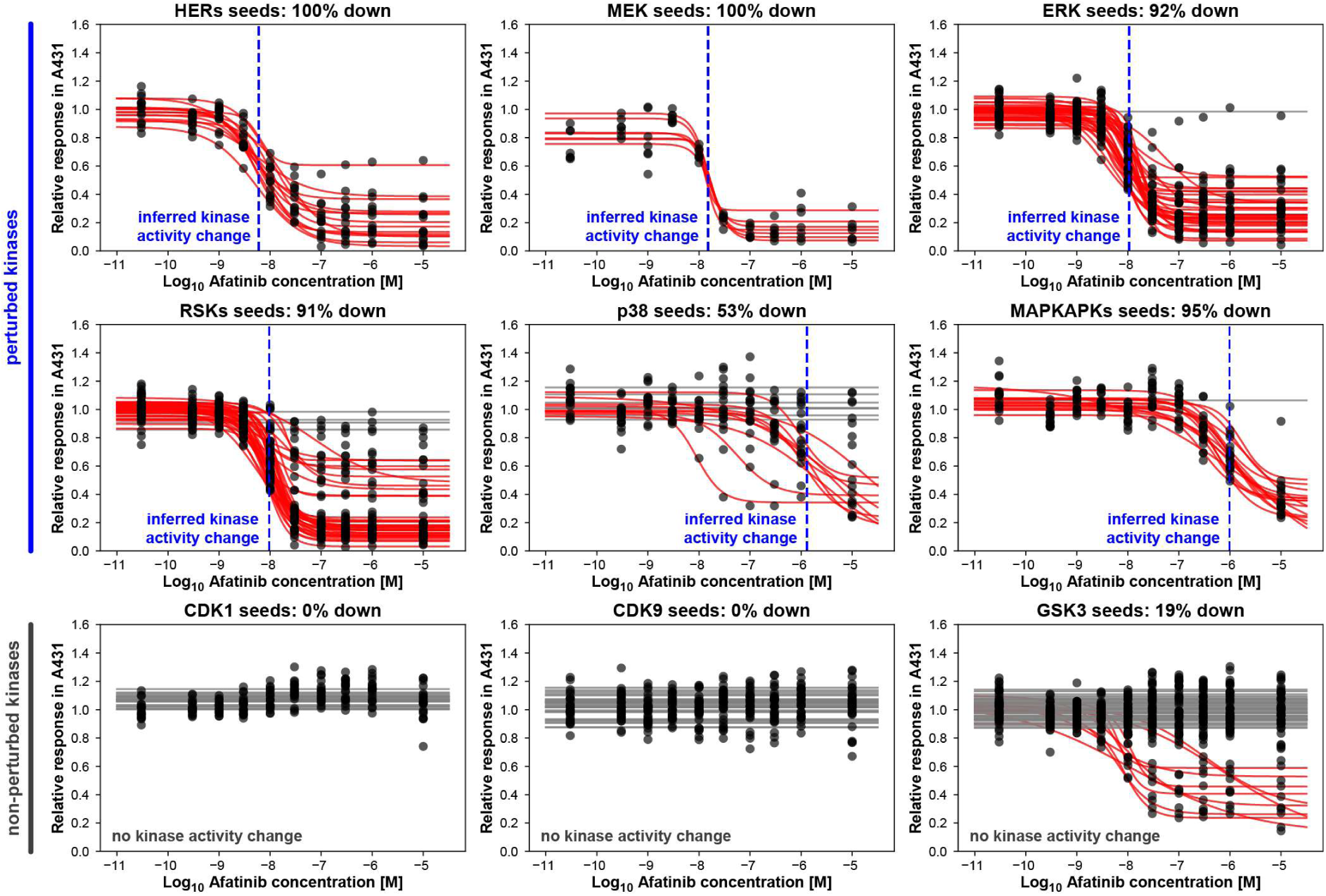
Inference of kinase perturbing concentrations from seed peptidoforms for different kinase groups in Afatinib-A431. *Example of the estimation process of different kinase activity changes from seed peptidoform substrate spaces for Afatinib in A431. Seed peptidoform dose-response profiles were plotted for different kinase groups (each subplot for a different kinase). Each profile (measured dots + regressed model line) in each subplot is a measured peptidoform. Red peptidoforms indicate down regulations, and gray peptidoforms indicate “not” regulations. Curve classification was performed by CurveCurator. If more than 33% of classified seeds were significantly perturbed (up or down), the kinase group was also considered to be perturbed because enough evidence had been collected to conclude this. The inferred kinase perturbing concentration (inf_pEC_50_, blue dashed line) is the median pEC_50_ value of all significantly perturbed seed peptidoforms. Please note that we did not require seeds to be strictly dynamic across all cell lines. Thus, some seed peptidoforms for ERK and RSKs may appear as not-regulated in some cell lines. Please also note that GSK3 possesses some down-regulated seeds, because all GSK3 substrates need to be primed by a different upstream kinase. If the priming kinase was perturbed, then the seed peptidoform would also be perturbed, even though GSK3 itself didn’t change activity*.

**Figure S6:**
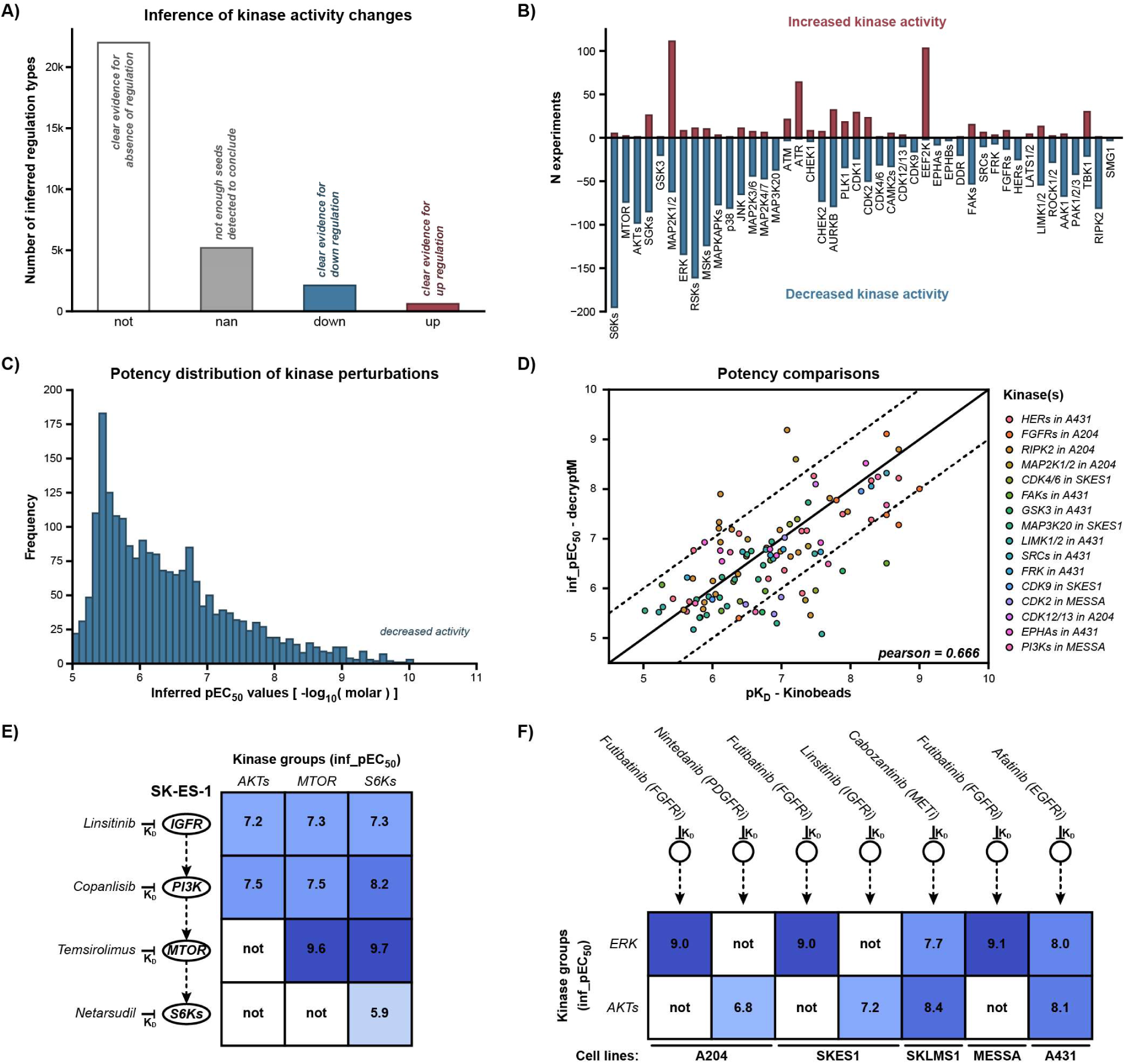
Different aspects of the kinase perturbation matrix. ***(A)*** *A summary statistic that counts the inferred kinase regulation types (not, down, up, undetermined) in the kinase perturbation matrix (see main figure 3D). **(B)** A summary statistic that counts how often a specific kinase in the kinase perturbation matrix was perturbed directly by the inhibitor or indirectly as a consequence of upstream perturbation. **(C)** Distribution of inf_pEC_50_ values of decreased kinase activities. **(D)** Correlation plot between inf_pEC_50_ values from decryptM and pK_D_ values from Kinobeads (Klaeger et.al). For each kinase, one reference cell line was chosen in which the kinase was clearly active, and then all inhibitors that have the kinase as a target were plotted. **(E)** Systematic inhibition along a linear pathway (rows) leads to stepwise upstream-downstream separation of kinase groups (columns). IGFRi and PI3Ki in SK-ES-1 co-inhibit the kinase groups AKTs, MTOR, and S6Ks. The inhibition of S6Ks only inhibits S6Ks but not upstream kinases AKTs or MTOR. **(F)** Systematic RTK inhibition across different cell lines (columns) shows different pathway connectivity in the different cell lines of ERK and AKTs (rows)*.

**Figure S7:**
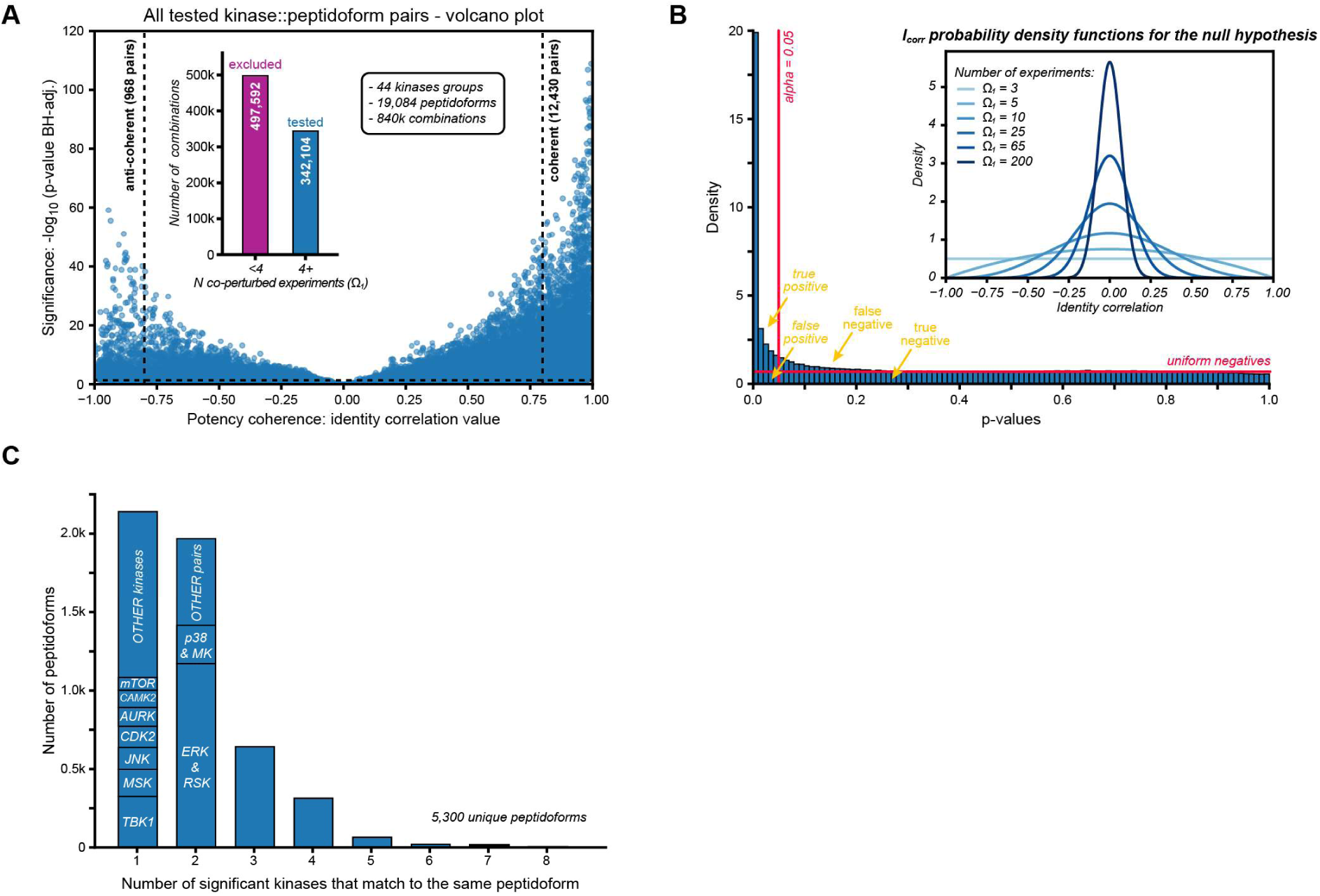
Global identity correlation analysis to identify potency-coherent kinase::substrate relationships. ***(A)*** *Volcano-plot representation of the potency coherence (x-axis: i_corr_) vs. the statistical significance (y-axis: p-value_adj._) of all kinase peptidoform pairs with four or more co-perturbations (Ω_1_>3). After Benjamini-Hochberg correction, statistical significance was cut at an p-value_adj._<0.05 (horizontal dashed line). An i_corr_>0.8 indicates high potency coherence between a kinase and a candidate substrate. An i_corr_<-0.8 indicates high potency anti-coherence with an opposite direction of regulation. **(B)** Experimental p-value distribution of all 342k tests from subpanel A. The shape of the experimental p-value distribution verifies well-calibrated p-values and the validity of the statistical approach. P-values were calculated using the single beta approximation for i_corr_ distributions that depend on the number of observations. Exemplary i_corr_ distributions of the null hypothesis H_0_ are depicted in the inlet for different numbers of data points. Ω_1_ refers to the number of co-perturbations. The number of dis- coherent experiments (Ω_2+3_) was 0 for these distributions. **(C)** Bar-plot of peptidoform counts mapping to multiple significant kinase::peptidoform pairs. Here, we excluded MAP2Ks form the analysis to simplify interpretation. There are often 1::1 relationships. The 2::1 relationships are dominated by ERK-RSKs and p38-MAPKAPKs (abbr. MK), mapping to the same peptidoforms, respectively. Higher-order relationships (4+::1) are rare*.

**Figure S8:**
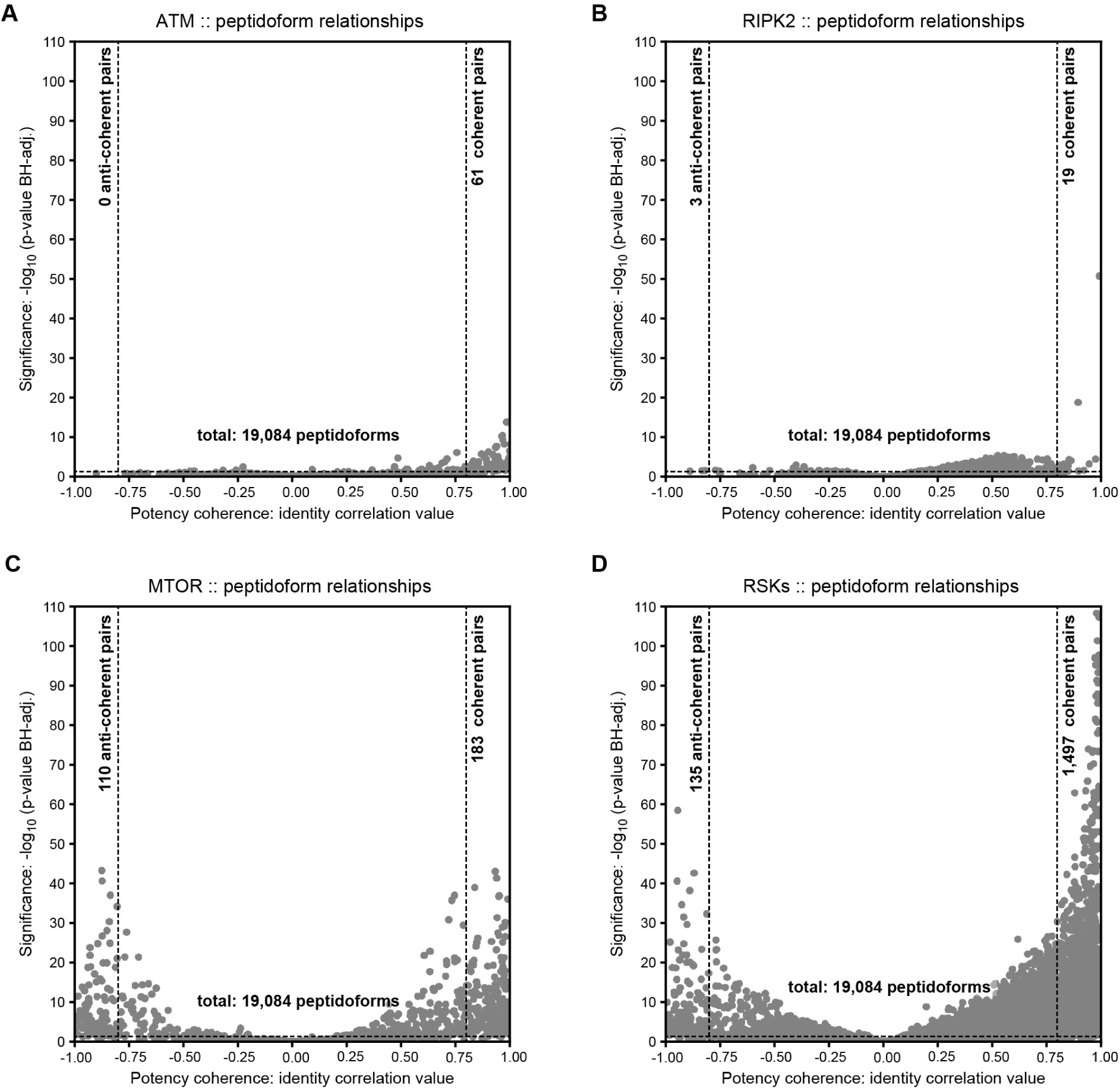
Potency coherence analysis for different kinases. *Volcano-plot visualizations (potency coherence vs. statistical significance) after calculating all kinase::peptidoform combinations for different kinases (**A:** ATM, **B:** RIPK2, **C:** MTOR, **D:** RSKs). P-values were Benjamini-Hochberg adjusted for the entire data set (not per kinase). The thresholds (dashed lines) are set at a ±0.8 identity correlation and a 0.05 adjusted p- value*.

**Figure S9:**
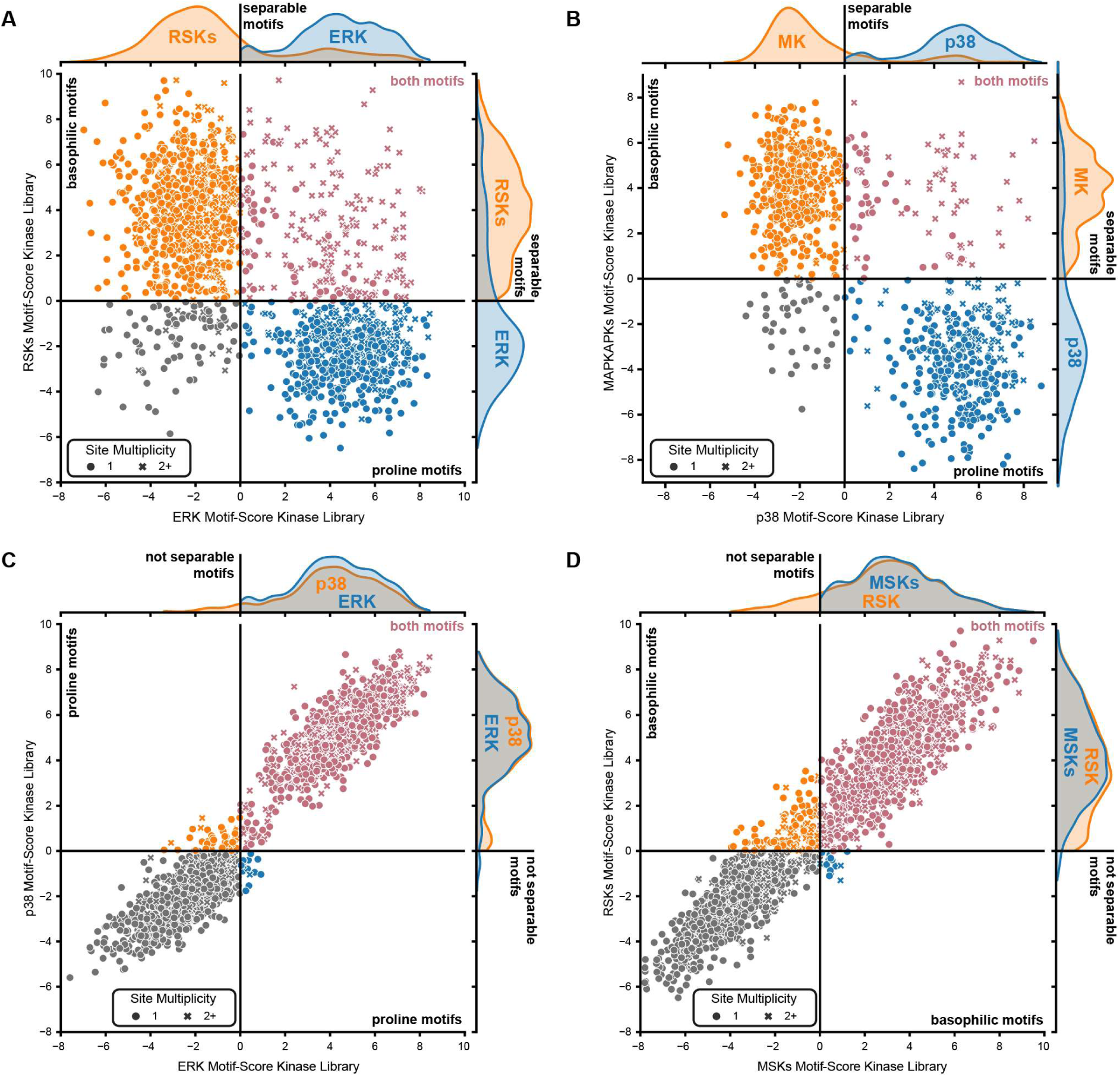
Separation of peptidoforms by kinase-library motif scores. *Scatter plots that relate how well a peptidoform contains a phospho-acceptor motif for a specific kinase group (x-axis and y-axis) based on the MIT Kinase Library. A dot indicates one phospho-site per peptidoform. A cross indicates two or more phospho-sites on the same peptidoform. A motif score (log_2_ odds) of 0 means that a motif is equally likely as random. Peptidoforms were colored by kinase plausibility based on filtering by motif scores. A kinase is not possible if motif scores are smaller than 0. If both motifs are plausible, the color is red. If no motif is plausible, the color is gray. **(A)** There is a clear separation by motifs between ERK and RSKs because they come from different evolutionary kinase families. **(B)** There is a clear separation by motifs between p38 and MAPKAPKs because they come from different evolutionary kinase families. **(C)** There is **<u>no</u>** separation by motifs between ERK and p38 because they come from evolutionarily close kinase families. They can be separated by potency coherence analysis (not visible here). **(D)** There is **<u>no</u>**separation by motifs between MSKs and RSKs because they come from evolutionarily close kinase families. They can be separated by potency coherence analysis (not visible here)*.

**Figure S10:**
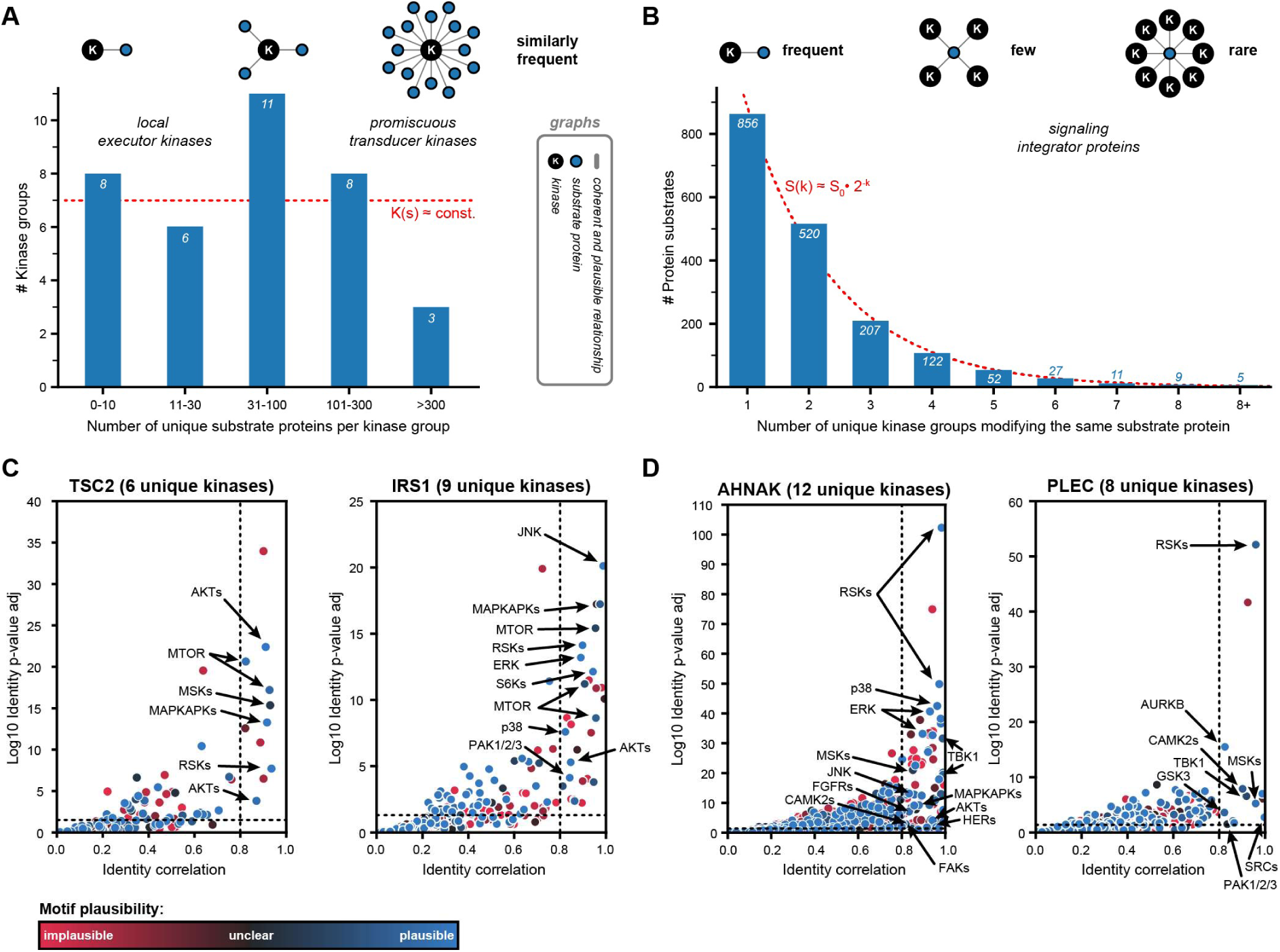
Kinase and substrate protein signaling hubs. ***(A)*** *A barplot visualizing the distribution of the number of unique substrate proteins that a kinase group has based on potency-coherent and motif-plausible KSRs. The bars were binned half-logarithmically. There is an equal amount of kinase groups with very few, medium, or many substrate proteins. This situation is illustrated by the three graphs. Each blue dot represents 10 substrates. **(B)** A barplot to visualize the distribution of the number of unique kinase groups per substrate protein based on potency-coherent and motif-plausible KSRs. The distribution follows an exponential function. This means that most proteins have only a few dynamic kinase relationships, and a few proteins have many dynamic kinase relationships. This situation is illustrated by the three graphs at the top. Each blue dot represents a substrate. **(C&D)** Volcano-plot visualization (potency coherence vs statistical significance) after calculating all peptidoform::kinase combinations for the proteins TSC2, IRS1, AHNAK, and PLEC. P-values were corrected for multiple testing using the Benjamini-Hochberg method across the entire data set. The thresholds are set at ±0.8 identity correlation and a 0.05 p- value_adj._. The dot color indicates the motif plausibility based on the MIT Kinase Library score (implausible: red, unclear: dark, plausible: blue). Exemplary kinases were highlighted for the substrate proteins. Every blue dot in the right upper box represents a potency-coherent and motif-plausible KSR*.

**Figure S11:**
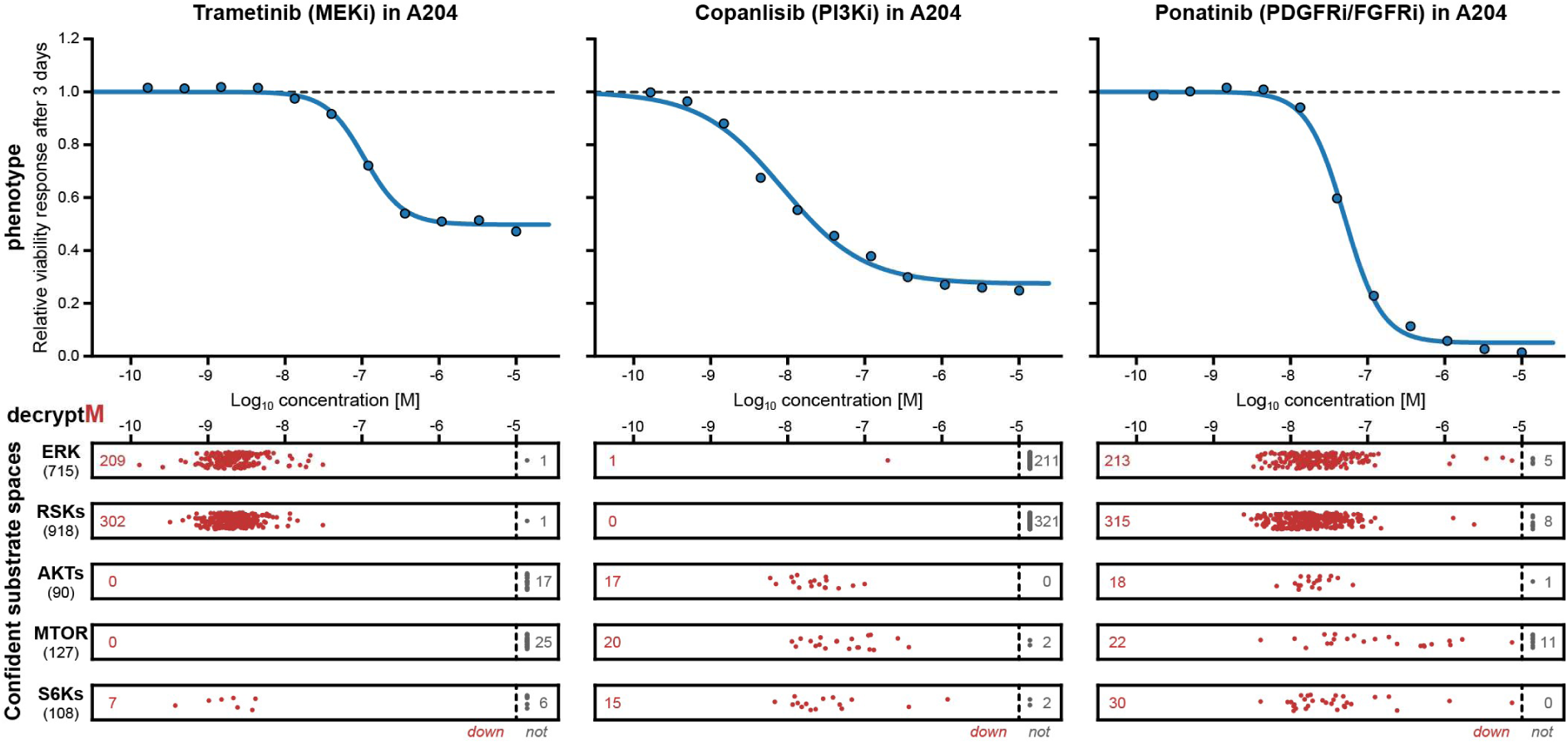
DecryptM and confident substrate spaces reveal the concomitant drug-induced proteoform changes that contribute to the phenotype. *Example for integration of cell viability data (blue dose-response curves; Lee et al.) with decryptM data (red swarm plot; this study) through the same and shared drug potency dimension. The decryptM data were filtered for confident KSRs and plotted by kinase groups (the size of substrate space is shown below each kinase group name on the left). Each red dot shows the pEC_50_ of the down-regulated peptidoform dose-response curve. Each gray dot depicts not-regulated peptidoforms. The number of down- or not-regulated peptidoforms is also stated in each box. This type of analysis builds the basis for main figure 5A*.

**Figure S12:**
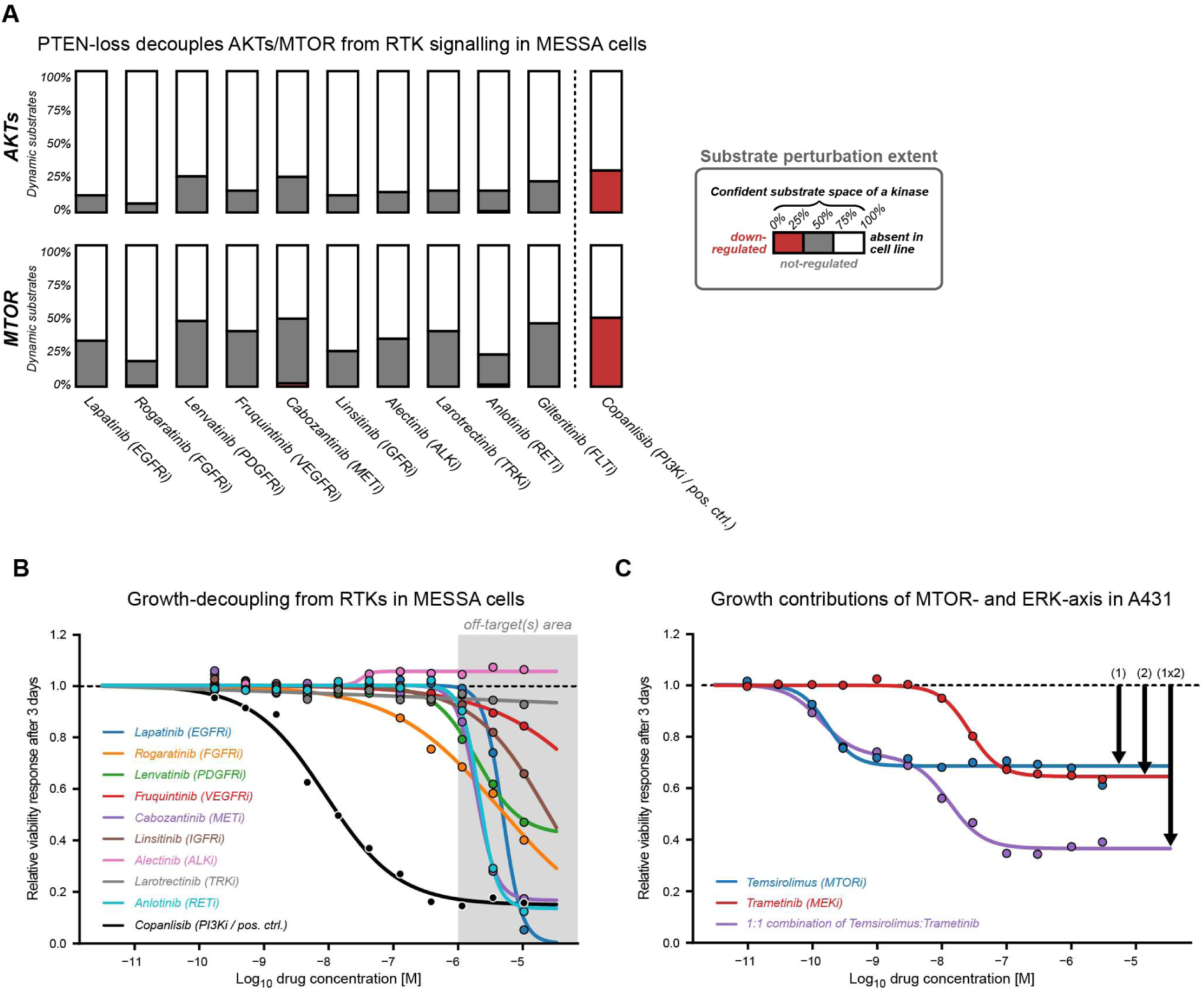
Signaling and growth contributions in MESSA and A431 cells. ***(A)*** *Substrate perturbation extent analysis based on decryptM data for different RTK inhibitors in MESSA. 100% refers to the complete potency-coherent and motif-plausible substrate space for a kinase as defined in this study. The color indicates the down-regulated (red), not-regulated (gray), or absent/unclear (white) proportion for a decryptM experiment. The potent, designated target is specified after the drug name. While Copanlisib (PI3Ki) could regulate all observed substrates of AKTs or MTOR, no RTK inhibitor could (all observed substrates were not-regulated). **(B)** Viability assay (Alamar blue) after 3 days of drug treatment in MESSA cells. The response was fitted with a 4-parameter sigmoid model. The different drugs were color-coded. Except for Rogaratinib (which inhibits growth via the ERK axis), no potent change in viability was observed. A grey area above 1 µM indicates responses due to off-target effects, since clinical inhibitors act on their designated targets at much lower concentrations than 1 µM. **(C)** Viability assay (Alamar blue) after 3 days of drug treatment in A431 cells. The response was fitted with a 4-parameter sigmoid model. For the combination treatment, Trametinib and Temsirolimus were mixed in a 1:1 ratio. This means that the x-axis values are correct for both inhibitors. The drug-combination dose-response curve was expectedly biphasic and was fitted with a 7-parameter double sigmoidal model to capture both phases. The potencies of the single treatments align with those of the double treatments*.

**Figure S13:**
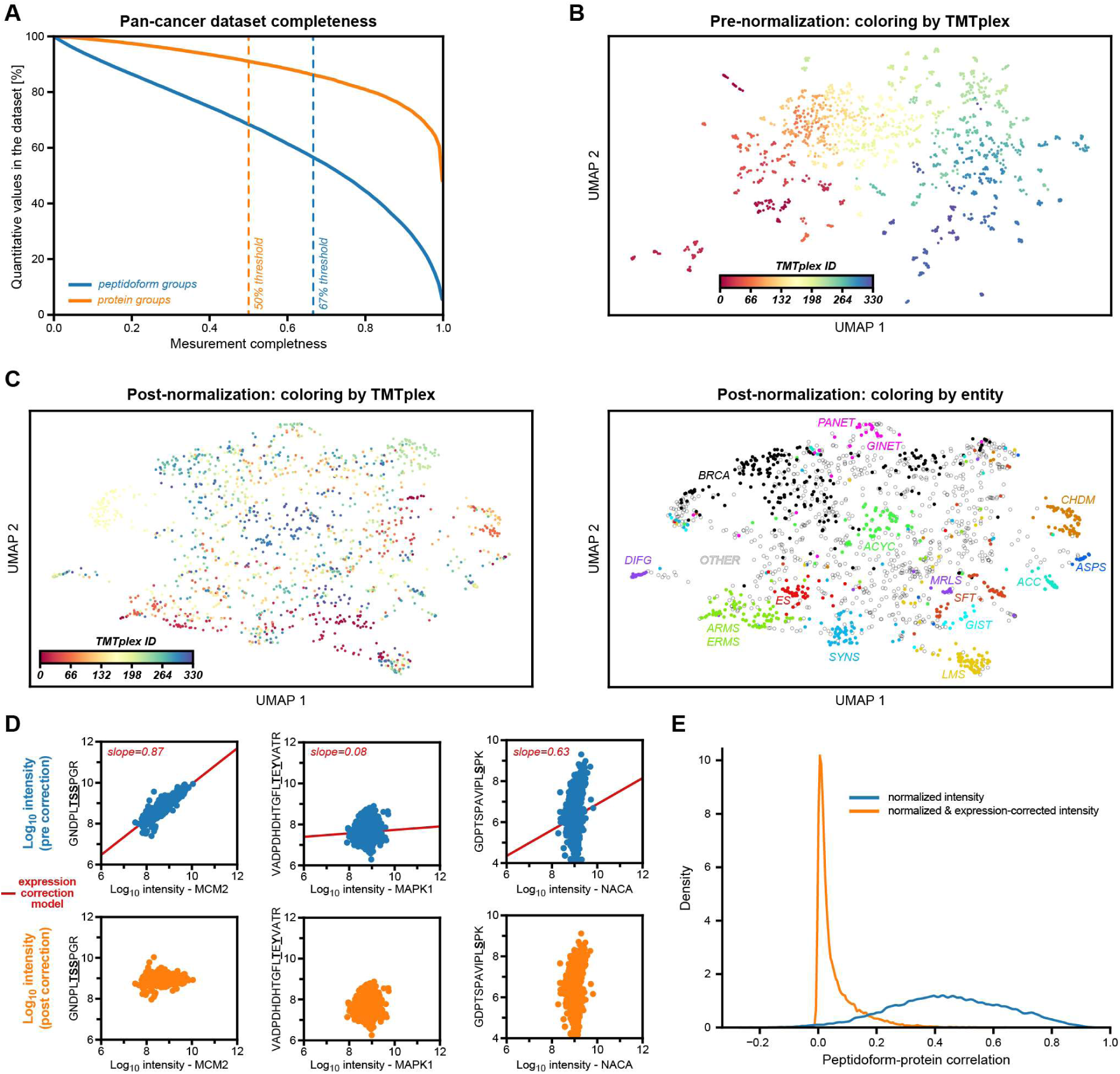
Patient data normalization procedure. ***(A)*** *Relationship between data completeness (detected in x% of patients) and retained quantitative values (relative to all values in the data matrix if a completeness threshold were applied) to see the consequence of data filtering. For the full proteome, we applied a 50% completeness threshold. For the phosphoproteome, we applied a 67% completeness filter*. ***(B)*** *Pre-batch-correction UMAP of the phosphoproteome, where each data point is a patient and the color indicates the TMT batch ID. The clustering is driven by TMT-plexing. **(C)** After the bridge-channel normalization, the TMT batch effect is gone, and the UMAP of the phosphoproteome clusters by cancer entity (= protein expression pattern in the phosphoproteome). Left and right are the same UMAPs, one colored by TMT batch and the other by selected entities. **(D)** Example cases to show different relationships between a peptidoform and the corresponding protein baseline expression. The left panel exemplifies a strong expression effect between the peptidoform and the protein. The middle panel is an example of weak dependence, and the right is an example of a high phospho dynamic range but a low protein dynamic range. The red line indicates the fitted log-log linear relationship (restricted to 0 <= slope <= 1) that was used to remove the expression effect in the phosphoproteome. The top row shows the abundance expression relationship before correction, and the bottom row shows the relationship after expression correction. **(E)** Distribution plot before and after correction that verifies that there is, in general, a fair amount of correlation between the peptidoform abundance and the protein expression. After applying the expression correction with the restricted model, this relationship is mostly gone*.

**Supplementary Figure 14.**
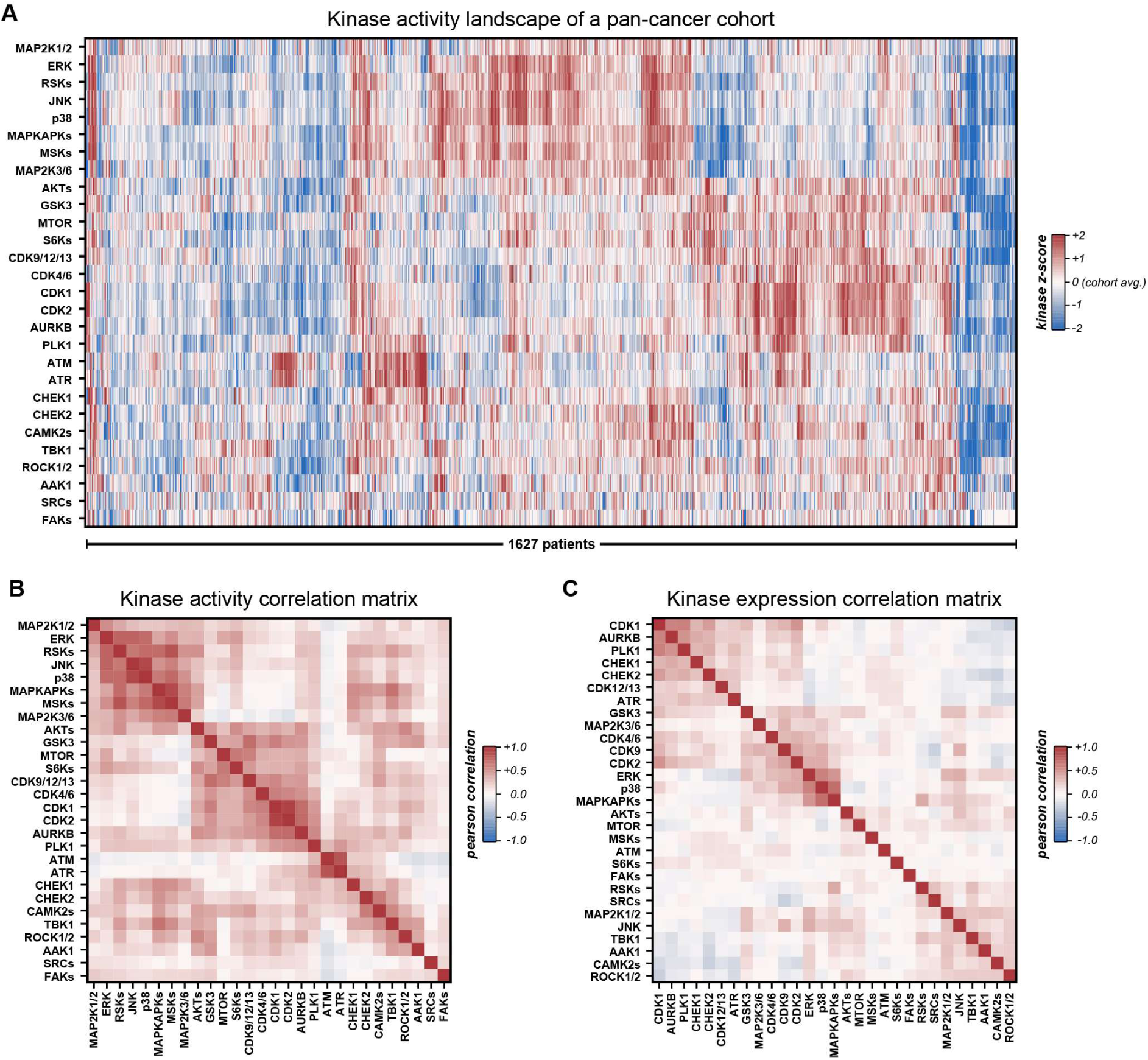
: Pan-cancer patient kinase activities. ***(A)*** *The kinase activity landscape for 28 kinase groups in the pan-cancer cohort that could be tracked with confident KSRs. Each row represents a kinase group, and each column corresponds to a patient. The color indicates the kinase activity z-score, where more red indicates greater activity and more blue indicates lower activity relative to the pan-cancer cohort (1,627 patients). **(B)** The kinase activity correlation matrix identifies kinases that have similar activity patterns across all patients in the cohort. The more red, the higher the positive correlation. **(C)** Same as in B, but here the protein expression is correlated. The more red, the higher the positive correlation. Most kinases have unique expression patterns and show low correlation to other kinases*.

## Methods

### 1. Cell culture

Cell lines used in this study were obtained from ATCC, Hölzel, or DSMZ. Cell lines were grown in media recommended by the manufacturers at 37 °C and 5% CO_2_, supplemented with FBS (PANbiotech: #P30- 3031). A204 (DSMZ: #ACC-250, McCoy’s 5a medium, 10%_v/v_ FBS), SK-ES-1 (DSMZ: #ACC 518, McCoy’s 5a medium, 15%_v/v_ FBS), SK-LMS-1 (Hölzel: #CLS300125, DMEM:F12 medium, 2 mM L-glutamine, 5%_v/v_ FBS), MES-SA (ATCC: #CRL-1976, McCoy’s 5a medium, 10%_v/v_ FBS), A431 (ATCC: #CRL-1555, DMEM medium, 10%_v/v_ FBS).

While preculture was performed in dishes, cell expansion was carried out in a 250 mL stirrer flask (Bellco IBO) at a concentration of 3 mg/mL Cytodex 1 (Cytiva) microcarriers until the cell density reached a target value of 1.0×10e6 cells/mL. Washing and conditioning of microcarriers were performed according to the manufacturer’s protocols. The cell seeding density was adjusted to 1.0×10e5 cells/mL. Half of the reactor medium was replenished when the glucose concentration decreased by more than 50% relative to the culture’s starting condition. 10 mM HEPES was supplemented in the end phase to stabilize the culture. Stirring speeds increased from 25 to 30 rpm during the 5- to 10-day expansion phase.

### 2. Drug treatments

The choice of compounds focused on the clinically most established kinase inhibitors that are currently either approved or in clinical phase III. The compounds were purchased from SelleckChem, MedChemExpress, or Hycultec. As manufacturers noted, the compounds were dissolved in DMSO, except for compounds that were not soluble in DMSO, which were instead solubilized in EtOH or H_2_O. Stock concentrations of 10 mM were stored at -80 °C. Drugs with solubility below 10 mM were adjusted to 3 mM or 1 mM instead. Drug-target affinity lists were obtained from previously published Kinobead results.

Before the drug treatment, 1 mL of the cell suspension was distributed to 2 mL deep-well filter plates (Porvair, #240002, 36 µm PE-fritted) and diluted with 0.7 mL conditioned medium of the cell expansion phase. The drug-working dilutions and the DMSO control were added to the wells and incubated on a plate rotator at 37 °C for 100 min (final concentrations: control, 0.03, 0.3, 1, 3, 10, 30, 100, 300, 1000, 10000 nM). After the treatment, cells were washed with 1 mL of PBS and then lysed in 150 μL of lysis buffer (2% SDS in 10 mM Tris-HCl, pH 7.6, 97 °C). Lysate was collected in a 96-well plate by centrifugation (1000 g, 2 min, RT) and stored at -20 °C until further use.

### 3. Sample preparation

The protein yield was determined using the Thermo Pierce BCA protein assay. All steps were performed according to the manufacturer’s protocol implemented on the Bravo Agilent pipetting system.

100 μg protein lysate per treatment condition was cleaned by protein aggregation capture on a Bravo Agilent pipetting system following the suggestions of the SP3 Protocol (*64*). Briefly, 100 μg protein lysate was mixed with 750 μg beads (1:1 mix of washed magnetic Sera-MagTM A & B; Cytiva) for 10 min at 1000 rpm RT. All volumes were adjusted to 120 μL using 10 mM Tris-HCl (pH 7.6). Proteins were precipitated by adding 100%_v/v_ ethanol to a final concentration of 70%_v/v_ and shaking for 10 min at 1250 rpm on the Bravo shaking unit. The supernatant was removed on the magnet. The lysate was washed three times with 80%_v/v_ ethanol, with 5-minute shaking intervals off the magnet and supernatant removal on the magnet. Finally, beads were washed with 100%_v/v_ ACN to remove residual ethanol. Proteins were reduced and alkylated in 100 μL RA buffer (100 mM EPPS/NaOH, pH 8.5, 50 mM CAA, 10 mM TCEP) for 1h at 37 °C and 1200 rpm. Next, 2 μg of Trypsin was added, and digestion occurred overnight at 37 °C and 1000 rpm. Then, peptides were recovered on the Bravo magnet. Beads were washed with 110 μL 2%_v/v_ TFA for 3 min at 1250 rpm on the Bravo shaking unit. The wash supernatant was pooled with the recovered peptides. To further remove magnetic beads from the digest, the recovery plate was incubated on a magnetic rack for 1 h at 4 °C.

The collected digest was desalted using HLB desalting plates (10 mg, N-Vinylpyrrolidone-Divinylbenzol (NVP–DVB), porous particles, 30 μm, Macherey-Nagel). Plates were cleaned with 400 μL 100%_v/v_ iPrOH, 100%_v/v_ ACN, and solvent B (0.1%_v/v_ TFA, 70%_v/v_ ACN), and then equilibrated with 1000 μL solvent A (0.1%_v/v_ TFA). The digest was loaded by gravity. After washing with 1000 μL solvent A, peptides were eluted by gravity in 200 μL solvent B. The cleaned peptides were dried down in the speed-vac and stored at -20 °C.

### 4. Dose-multiplexing with TMT

The TMT-labeling followed the down-scaled TMT protocol (*65*) with slight modifications to improve throughput and robustness. The dried and cleaned peptides were reconstituted in 20 μL of 100 mM EPPS*NaOH (pH 8.5) buffer. TMT-11plex reagent (Thermo Scientific, LOT: UG291261) was reconstituted in water-free ACN to a working concentration of 20 μg/μl. 5 μL of this TMT reagent solution was transferred to the peptides. The reaction was incubated on the thermoshaker (23 °C, 400 rpm) for one hour and quenched with 0.25%_v/v_ hydroxylamine (NH_2_OH) afterward. The TMT channels were then pooled together and acidified with TFA to a final concentration of 1%_v/v_. The reaction wells were washed with 25 μL washing solution (30%_v/v_ ACN, 2.5%_v/v_ TFA) and added to the TMT pool. The TMT pools were dried down in the speed-vac and stored at -20 °C.

The 1.1 mg TMT-pooled peptides were cleaned by solid-phase extraction on 50 mg C18 Sep-PAK cartridges (Waters). The TMT peptides were washed with 0.1%_v/v_ TFA. TMT peptide elution was achieved by 0.1%_v/v_ TFA in 50%_v/v_ ACN. TMT peptides were dried down in the speed-vac and stored at -20 °C.

### 5. Phosphoproteome enrichment

Phosphorylated peptides were enriched by immobilized metal ion affinity chromatography (IMAC) on a ProPac 0.5 mL Fe(III)-loaded NTA column (Thermo Scientific). The column was connected to an Aekta Pure LC system (Cytiva). The dried TMT-pooled peptides were reconstituted in 175 μL solvent A (0.07%_v/v_ TFA, 30%_v/v_ ACN) and loaded onto the column at a flow rate of 0.2 mL/min. Non-phosphorylated peptides could not bind to the column and were collected by a fraction collector. Then, 5 mL of solvent A was used to wash the bound peptide at a flow rate of 3 mL/min, followed by a step elution with 42.5%_v/v_ solvent B (0.315%_v/v_ NH_4_OH in H_2_O) at a flow rate of 0.55 mL/min. The 900 μL phospho-fraction was collected and acidified with TFA to a final concentration of 1%_v/v_ TFA. The flow- through fraction and the phospho fraction were dried down in the speed-vac and stored at -20 °C.

### 6. bRP fractionation

The phosphorylated, TMT-labeled peptides were bRP-fractionated on RPS cartridge tips (5 μL PS-DVB, Agilent) into ten fractions using the Agilent AssayMAP Bravo pipetting system. The RPS cartridges were washed and equilibrated according to the manufacturer’s protocol. The peptides were reconstituted in 100 μL of 1%_v/v_ FA and loaded onto the cartridges. The cartridge flow-through was reapplied. The peptides were then washed once with 0.1%_v/v_ FA, followed by a pH change with 50 μL 25 mM Ammonium formate (pH 10). Peptides were fractionated under these basic conditions with increasing ACN concentrations (5%_v/v_, 7.5%_v/v_, 10%_v/v_, 12.5%_v/v_, 15%_v/v_, 17.5%_v/v_, 20%_v/v_, 25%_v/v_, 30%_v/v_, 70%_v/v_). The ten elution steps were pooled back to 6 fractions in an n-with-(n+6) fashion. All fractions were acidified with FA to a final concentration of 1%_v/v_. The fractionated phospho-TMT peptides were dried down in the speed-vac and stored at -20 °C until MS measurement.

### 7. LC-MS measurement

Dried samples were reconstituted in 12 μL 0.1%_v/v_ FA, and 10 μL were injected per fraction. Peptides were measured with an Eclipse Tribrid mass spectrometer (Thermo Scientific) that was coupled to a Dionex UltiMate 3000 RSLCnano System (Thermo Scientific). After injection, the sample was transferred to a trap column (75 μm × 2 cm) packed with 5 μm C18 resin (Reprosil PUR AQ - Dr. Maisch). Peptides were washed with the trap washing solvent (0.1%_v/v_ FA, 5 μl/min, 10 min) before conveying them to an analytical column (75 μm × 48 cm) that was packed with 3 μm C18 resin (Reprosil PUR AQ - Dr. Maisch). Separation was performed on an 80 min gradient with a flow rate of 300 nl/min starting from 4%_v/v_ B, followed by the first linear phase to 22.5%_v/v_ B in 65 min, followed by the second linear phase to 32%_v/v_ B in 15 min. The system was finally washed with 80%_v/v_ B for 2 min and re-equilibrated at 2%_v/v_ B. Solvent A consisted of 0.1%_v/v_ FA and 5%_v/v_ DMSO in H_2_O. Solvent B consisted of 0.1%_v/v_ FA and 5%_v/v_ DMSO in ACN.

The MS was operated in a sensitive, data-dependent MS^3^-mode. Peptides were ionized using a nano source with 2.1 kV spray voltage. Every 3 s, a full-scan (MS^1^) was recorded from 360 to 1800 m/z at a resolution of 60,000 in the Orbitrap in profile mode. The MS^1^ AGC target was set to 4 × 1e5, and the maxIT was set to 50 ms. Based on the full scans, precursors were targeted for MS^2^ scans if the charge state was between z=2 and z=5, the isotope envelope was peptidic (MIPS), the intensity exceeded 5 × 1e4, and the precursor mass lay between 940 and 5400 Da. The MS^2^ quadrupole isolation window was set to 0.7 Th. Peptide fragmentation occurred in the linear ion trap by CID-targeting the precursor and the precursor-H_3_PO_4_ (NL = 97.977 Da) in parallel (multistage-activation) with a q-value of 0.25, 35% CE, and 10 ms activation time. The MS^2^ spectrum was acquired at 30,000 resolution and with an auto- scan range in the Orbitrap in centroid mode. The MS^2^ AGC target was set to 1.5 × 1e5 charges, and the maxIT was set to 60 ms. Precursors that have been targeted for fragmentation were excluded for 90 s for all possible charge stages. TMT reporter ions were measured in a consecutive MS^3^ scan based on the previous MS^2^ scan. Thus, a new batch of precursor ions was isolated with a charge-stage-dependent MS^3^ quadrupole isolation window of 1.2 Th (z=2), 0.9 Th (z=3), and 0.7 Th (z=4&5) to include the first isotope in the isolation while using the narrowest isolation window possible. The isolated precursor was then MSA-fragmented identically to the previous MS^2^ scan. The top 10 fragment ions of the MS^2^ scans were isolated in the ion trap in parallel (synchronous precursor selection; SPS). Only fragment ions were considered that were within a range of 400 to 2000 Th and lay outside the precursor exclusion range (precursor -65 Th, precursor + 5 Th). Additionally, isobaric tag loss exclusion properties were set to TMT reagent. The selected top 10 fragment ions were then HCD fragmented with an NCE of 55%. The MS^3^ spectrum was acquired at 50,000 resolution from 100 to 1000 Th in the Orbitrap in centroid mode. The MS^3^ AGC target was set to 2.5 × 1e5 charges, and the maxIT was set to 120 ms.

### 8. Peptidoform identification and quantification

Peptide identification and quantification were performed using an in-house pipeline that comprised various published tools to process ∼4000 90-minute raw files (total ∼3 TB) in the most efficient and consistent manner possible.

All raw files were directly parsed using the ThermoFisher RawFileReader (version 5.0.7.1) to obtain all available information for each spectrum, including explicitly TMT report information from the MS^3^ scans, such as raw signal, raw noise, and resolution. TMT intensities were corrected for isotopic impurities according to TMT-LOT UG291261 using numpy’s linalg.lstsq solver. Signal-to-noise ratios were calculated for each reporter as a quality metric.

MS^2^ spectra were converted to mzML files using MSConvert (version 3.0.20335)(*66*). MS^2^ spectra were then clustered using MaRaCluster (version 1.01.1)(*67*) for each cell line separately to identify similar MS^2^ fragmentation spectra within and across runs.

MaxQuant (version 1.6.17.0) with its built-in search engine Andromeda (68) was used to annotate MS^2^ spectra with sequence information from a human UniProt database (97,086 known isoforms from Nov. 2020) supplemented with common contaminants at 100% false discovery rate (FDR). Unless stated otherwise, default parameters were applied. Trypsin/P was set as the proteolytic enzyme, allowing up to three missed cleavage sites. Carbamidomethylation of cysteine was set as a fixed modification. Oxidation of methionine, N-terminal protein acetylation, and phosphorylation on serine, threonine, and tyrosine residues were allowed as variable modifications. A fixed TMT-11plex modification was specified for lysine and peptide N-termini. Precursor tolerance was set to ±5.0 ppm and fragment ion tolerance to ±20 ppm (FTMS). All PSMs were additionally re-scored using pyAscore (version 1.0.1) (*69*). Percolator (version 3.02.1) (*70*) was used to adjust FDR to 1% at the peptidoform level for each cell line, using the following input features: Andromeda delta score, pyAscore peptide score, precursor mass, absolute precursor mass error, number of charges, and number of phosphorylations. Confident peptidoform identifications (percolator PEP < 0.1%) were ID-harmonized and SIMSI-transferred (*71*) within a MaRaCluster p20 cluster as long as the transferred pyAscore peptide score > 120. The most PEP-confident MS^2^-MS^3^ measurement pair was used as the reference for peptidoform identification and quantification in further analysis.

### 9. Normalization, curve fitting, curve classification

Each decryptM experiment, i.e a specific drug-cell line combination, was analyzed separately by CurveCurator (version 0.5.1) (*26*). First, TMT channels were normalized by column-wise log-median-centering, and zeros were imputed. Next, ratios were calculated relative to the control channel. Then, the four-parameter sigmoidal curve was regressed using the efficient multi-guess OLS algorithm, which yields, among other parameters, potency (pEC_50_ = -log_10_ EC_50_) and efficacy (log_2_ curve fold change) estimates for each peptidoform. While potency is a directly regressed parameter, efficacy is defined as the log_2_ ratio of the regressed model between the lowest and highest concentration.

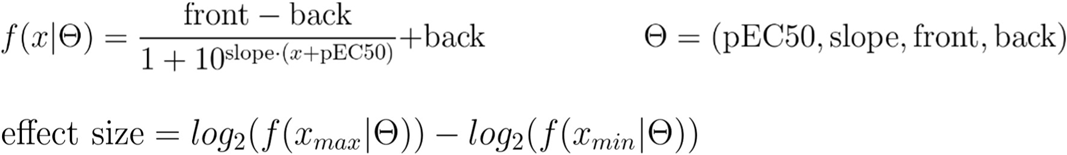

Curve relevance was defined by an alpha-limit of 5% and a fold change limit (fc_lim_) of ±0.45 for “up” and “down” curves. “not”-regulated curves had a mean model RMSE<0.1. An additional max pEC_50_ filter was set to 10 for relevant curves unless otherwise specified. All other curves that were neither “up”-, nor “down”-, nor “not”-classified remained unclassified and are not considered for downstream potency-based analyses.

### 10. Quality controls

Every TMT batch was subject to extensive quality control analysis, and some quality metrics are presented in figure S1. This included digestion efficiency, alkylation efficiency, TMT labeling efficiency, phospho-enrichment efficiency, and ID rate. Each TMT channel was also analyzed for overall intensity distributions, normalization factors, and systematic regression bias. The latter is measured by the median absolute deviation (MAD) across all peptides relative to the regressed model. A MAD <0.05 [ratio units] indicates excellent, 0.06 - 0.1 [ratio units] good, 0.11 - 0.15 [ratio units] poor, and >0.15 [ratio units] bad quality. TMT channels with bad MAD values were always excluded. TMT channels with poor MAD were manually evaluated to determine if they may be excluded from the analysis. If a channel was excluded, the experiment was refitted and classified with CurveCurator.

### 11. Dataset-wide aggregation of peptidoform dose-response curves

All analyses that require the integration of multiple decryptM perturbation experiments were performed on an aggregated data matrix. If multiple spectra were obtained for the same peptidoform in a single experiment, the spectrum with the lowest PEP was chosen. To further reduce missing values due to ambiguous localization, peptidoforms were grouped into locally delocalized peptidoform modification groups. The most prevalent peptidoform within the group was defined as the representative modified sequence. If there were conflicting observations, e.g., down- and up-regulated members for the same modification group, the group was set to NaN.

### 12. Qualitative perturbation comparisons

For pairwise comparisons of perturbed peptidoforms between two decryptM experiments, E1 and E2, only classified responses (“up”, “down”, “not”) were used. The similarity degree (Jaccard index) and overlap degree (Szymkiewicz–Simpson coefficient) were modified such that they separate “up” and “down” regulations and ignore “co-not” classifications. This gives the following formulas:

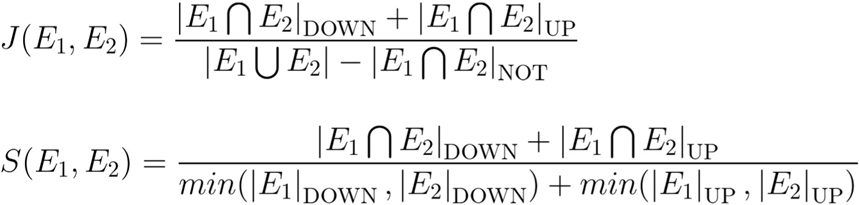

The clustering map was produced using sklearn’s (version 1.6.1) pairwise_distances function combined with AgglomerativeClustering. Cityblock was used as a distance metric, and clustering was performed using average linkage. Scipy’s (version 1.14.1) optimal_leaf_ordering algorithm was used to arrange the clusters in an optimal way.

### 13. Peptidoform annotations

Peptidoforms were annotated using the open-source p-site annotation package (version 0.5.3). Prior kinase substrate relationships were obtained from PhosphositePlus (Nov. 2022) (*5*). Annotations were merged with a localization uncertainty of ±2 to the representative modified sequence. If multiple annotations or annotation sites on the same peptidoform existed, they were all merged and considered.

Kinase motif scores were calculated based on the Kinase Library (*18, 19*) for each possible flanking sequence with and without additional modifications. A kinase motif score is essentially a log_2_-odds score, which is based on the multiplied amino acid preference of a kinase over positions in the flanking sequence. Thus, a value of 0 indicates that a motif for a kinase is as favorable as random. A value >0 indicates a preferred motif, and a value <0 indicates a disfavored motif. These raw motif scores were then sigmoid-transformed to obtain values in the range [-1, 1]. These transformed motifs were used as standardized input features for dimension-reduction techniques. For peptidoforms with multiple phosphorylation sites, the best of the multiple possible motifs was used. If a kinase group was evaluated, the best possible member of the group was used. If a motif type was evaluated, the best possible member of the same motif family was used. Kinase-peptidoform pairs with motif-scores >0 are called motif-plausible.

### 14. Kinase perturbation inference in decryptM

Kinase genes were grouped into functional kinase protein groups. Functional kinase groups have manually curated “seed sites” that we mapped to “seed peptidoforms”, which are indicative of kinase activity. The goal of the curation was to create a small but potency-reliable group of peptidoforms to estimate the potency of a kinase perturbation. The curation was mainly based on selective inhibitors (according to Kinobeads results (*30*)), from which decryptM profiles could be manually interpreted. The sites were then tested to follow the inhibitor’s expected potency patterns either through direct inhibition or indirect signal transduction of the perturbation, as observed with more complex drugs. A seed peptidoform must also have a plausible motif and ideally is already known from the literature. The latter, however, is not a strict prerequisite. A seed peptidoform should also be unambiguous and can therefore only map to one kinase group for the set of so far tested 5 cell lines. Given the inhibitors and cell lines used in this study, we obtained seed sites for 44 kinase groups.

To systematically assess if a kinase has changed activity and, if so, at which potency, we mapped the corresponding seed sites using the p-site annotation package. If >1/3 of mappable seed annotations of a kinase have changed upon treatment in a consistent manner, i.e. “up” or “down”, the kinase is deemed up- or down-regulated, respectively. Unclear and missing values were excluded from the analysis. The inferred potency is the median potency of all mapped down- or up-classified peptidoforms. Downregulation is denoted by a (-) sign, and upregulation by a (+) sign throughout this study. If no activity change was detected because most seed peptidoforms were “not”-classified, the kinase potency was set to 0. If all seed peptidoforms were unclear or missing for a kinase, the kinase potency was set to NaN because neither perturbation nor absence of perturbation could be concluded.

### 15. Potency coherence analysis

The first step of the potency coherence analysis is to calculate transformed EC_50_ values (tEC_50_). The pEC_50_ value is defined as the negative decadic logarithm of the effective concentration. The pMD value is defined as the negative decadic logarithm of the highest applied dose. The sign function maps the curve regulation type of the dose-response model “down” (-), “not” (0), and “up” (+). Please note that pEC_50_ values that are not “up”- or “down”-regulated (CurveCurator classification) have no biological interpretation and were excluded from the analysis or set to 0 in the case of “not”-classifications. In summary, the tEC_50_ value is defined as the signed effective concentration relative to the highest applied dose. To illustrate the concept see below.

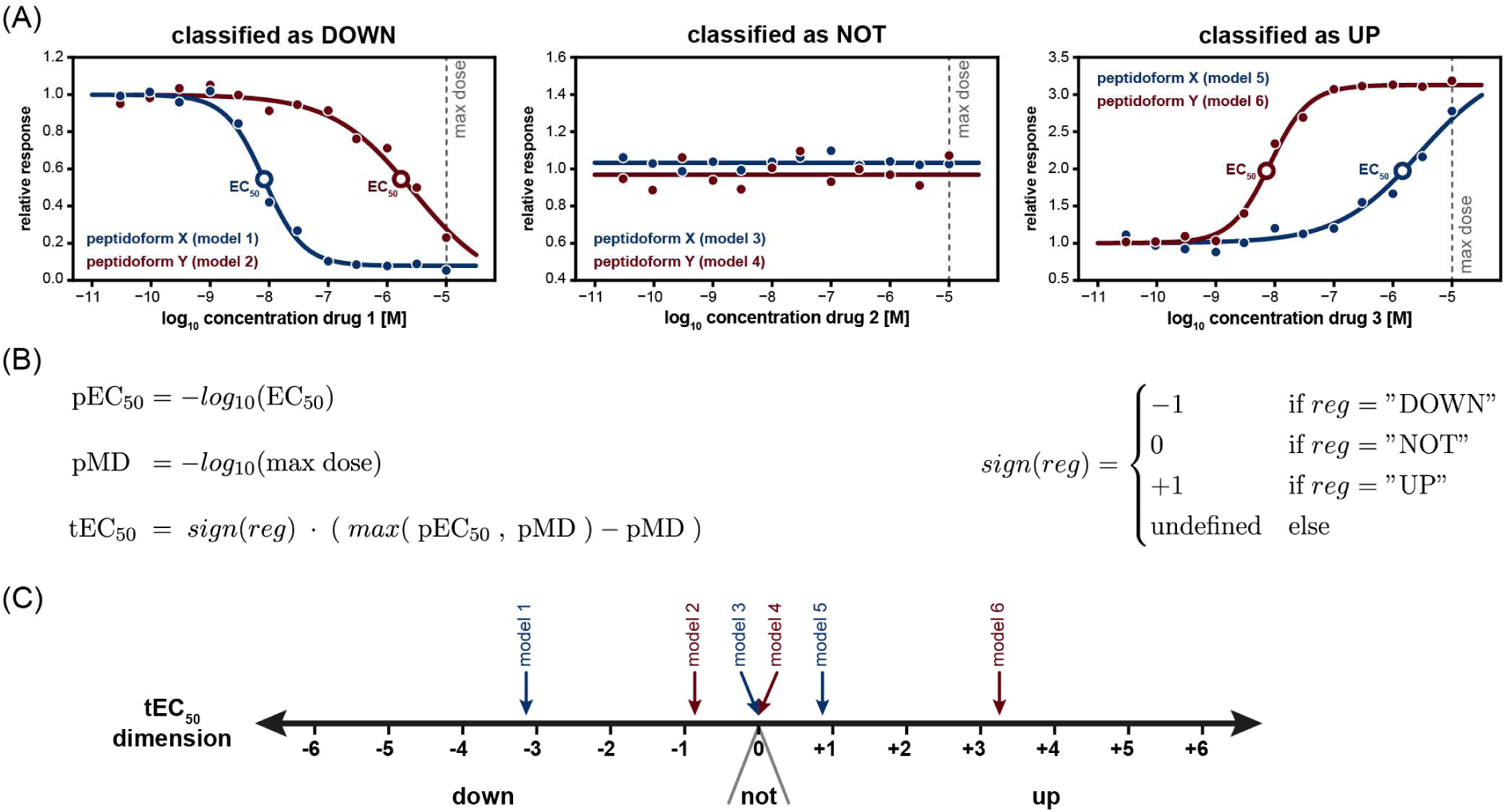

To quantify the degree of potency coherence between two transformed potency (tEC_50_) arrays X and Y, which can be visualized as a 2D potency coherence map, we defined the identity correlation value (I_corr_).

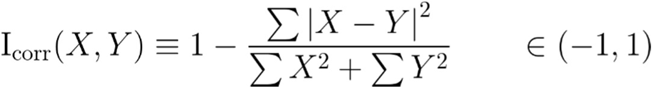

Given the number of valid potency observations N encoded in the configuration 𝛀, we can calculate a p-value for a specific identity correlation value using a single Beta distribution U with parameters alpha = beta = f(𝛀), shifted by -1 and scaled by 2.

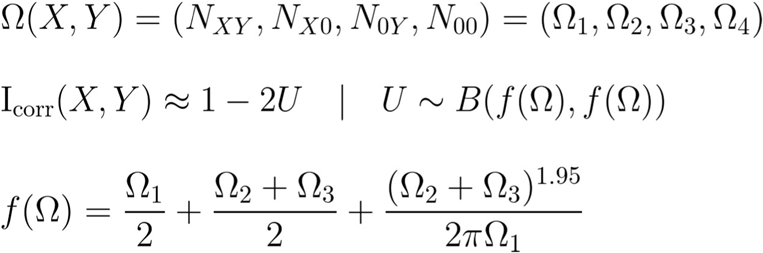

To estimate the proportion of potency-coherent peptidoforms in a set, all pairwise combinations were evaluated using the identity correlation value. The square root of the proportion of pairs with i_corr_>0.8 yields an upper-bound estimate. This analysis was applied to estimate the potency consistency of PhosphoSitePlus annotations. To be conservative, we considered only decryptM-perturbable substrate peptidoforms and used only pairs with (𝛀_1_+𝛀_2_+𝛀_3_)>=3 to build the combinatorial distributions.

### 16. High-throughput mining of kinase::peptidoform relationships

Kinase::peptidoform relationships are indicated by a “::” symbol and are categorized by different evidence levels (*hypothesis < candidate < confident < true*). All possible combinations are initially “*hypothesis”* relationships, which are systematically evaluated. Some of these hypotheses become “*candidate”* kinase::peptidoform relationships if they exhibit significant and high potency coherence (i.e., an identity correlation value greater than 0.80, a multiple-testing-adjusted p-value using the Benjamini-Hochberg method (*72*) smaller than 0.05, and a maximal mean potency difference less than 0.60). “*Confident*” kinase::peptidoform relationships additionally require a plausible phosphorylation motif (log_2_ motif odds score greater than 0.00 from the MIT Kinase Library; see above). Only if all theoretical alternative kinases can be experimentally disproven and one confident kinase::peptidoform relationship remains, do we obtain a “*true*” and “*unique*” relationship.

All kinase::peptidoform relationships can be visualized using potency-coherence volcano plots (coherence vs. significance). The color of the dots typically indicates the motif plausibility. These plots are available at ProteomicsDB (proteomicsdb.org/analytics/KSR).

### 17. Substrate perturbation extent analysis of kinases

The *“substrate space”* of a kinase group is defined as the complete set of peptidoforms that are both potency-coherent and motif-plausible for that kinase group across all cell lines. In a specific cell line, however, this substrate space is rarely fully present. We define the “*substrate perturbation extent*” of a kinase as the cell-line-specific subset of substrate peptidoforms that are both detected and responsive to drug treatments in a cell. This extent can be quantified either in absolute terms (the cardinality of the subset) or relative to the entire substrate space. To reduce the influence of missing values, multiple experiments can be merged for the analysis.

### 18. Kinase activity inference in clinical data

Clinical full- and phospho-proteomes were reprocessed from *Schneider et. al.* data set. We only considered QC-passed samples with >30% tumor cell content. Peptidoforms were locally delocalized (±2). The most prevalent peptidoform within the group was defined as the representative modified sequence. Peptidoform and protein groups were filtered to have >67% occurrence across all TMT batches. Different TMT batches were row-wise normalized using two bridge channels per batch (the 10th and 11th), which are a mix of five cell lines. Since protein expression also affects peptidoform abundance, we removed the protein expression contribution for each peptidoform by fitting a log-log-linear model with a bounded slope between 0 and 1, and subtracted the regressed linear protein expression effect from the phosphosite abundance. This expression-free abundance was then location-shifted to zero to remove the peptidoform-specific MS-response factor, which yields a lognormal distribution N(0, σ) for each peptidoform. The spread of sigma captures the peptidoform-specific dynamics across the cohort. Not standardizing sigma to 1 has the benefit of weighting highly dynamic sites more than less dynamic sites. Peptidoforms were annotated with dynamic, confident (motif- plausible and potency-coherent) kinase::substrate relationships from this study. For each kinase, the log-normal expression-free abundances of all substrate peptidoforms were summed. To compare the cohort distribution for different kinases with varying substrate set sizes, the sums were converted to z-scores. A kinase with a 0 z-score indicates pan-cohort average kinase activity. Negative values indicate lower-than-average activity, and positive values indicate higher-than-average activity relative to the pan-cancer cohort.

